# Unified control of temporal and spatial scales of sensorimotor behavior through neuromodulation of short-term synaptic plasticity

**DOI:** 10.1101/2022.10.28.514318

**Authors:** Shanglin Zhou, Dean V. Buonomano

## Abstract

Neuromodulators such as dopamine have been shown to modulate short-term synaptic plasticity (STP). Here we propose that the neuromodulation of STP provides a general mechanism to scale neural dynamics and motor outputs in time and space. We trained RNNs that incorporated STP to produce complex motor trajectories—handwritten digits—with different temporal (speed) and spatial (size) scales. The learned dynamics underwent temporal and spatial scaling when higher synaptic release probabilities corresponded to higher speed/size. Neuromodulation of STP enhanced temporal or spatial generalization compared to weight modulation alone. The model accounted for the data of two experimental studies involving flexible sensorimotor timing. Our results address a long-standing debate regarding the role of dopamine in timing and predict novel mechanisms by which dopamine may slow down neural dynamics and thus slow “clock” speed.

## INTRODUCTION

A universal feature of motor behavior is the ability to flexibly adjust the temporal and spatial scale of motor outputs. In the temporal domain, it is possible to produce very similar motor output patterns at different speeds or overall durations (Cicchini et al., 2012; Hardy et al., 2018; Remington et al., 2018). For example, people can flexibly control the tempo of a musical piece or the duration it takes to sign their names by altering their writing speed. Analogously, in the spatial domain, we can also flexibly change the size of one’s handwriting depending on the writing surface area available (Harpaz et al., 2014; Rosenbaum, 2010). Similarly, in the sensory timing domain, the brain can not only distinguish between different intervals but the encoding of interval length can be flexibly modulated by a range of factors, including dopamine levels (Buhusi and Meck, 2002; Drew et al., 2003; Soares et al., 2016).

It is increasingly clear that motor control and its spatial and temporal flexibility, are in part governed by the neural dynamics of recurrent neural networks (Churchland et al., 2012; Crowe et al., 2014; Hennequin et al., 2014; Merchant et al., 2015; Saxena et al., 2022; Stroud et al., 2018; Vyas et al., 2020), suggesting that the neural dynamics of recurrent neural networks themselves may undergo transformations that underlie both temporal and spatial scaling. However, the neural circuit mechanisms underlying flexible temporal and spatial transformations remain largely unknown. Some neurocomputational models have demonstrated that it is possible to temporally scale RNN dynamics—that is, speed up and slow down the speed at which neural dynamics unfolds—by providing a “speed” input (Hardy et al., 2018; Remington et al., 2018; Saxena et al., 2022; Stroud et al., 2018; Wang et al., 2018) or adjusting the neural input-output gains(Lindén et al., 2022; Stroud et al., 2018). However, the mechanisms underlying spatial scaling of RNN dynamics, that is, the amplitude of the neural trajectories, remains mostly unaddressed, but see(Lindén et al., 2022).

Here we propose a novel and biologically inspired mechanism based on the neuromodulation of STP, to flexibly govern both the temporal and spatial scales of RNN dynamics and sensorimotor behaviors. STP refers to a universal form of use-dependent synaptic plasticity that operates on the subsecond time scale (Abbott and Regehr, 2004; Motanis et al., 2018; Zucker and Regehr, 2002). Despite its presence at almost all synapses in the brain, the computational functions of STP remain poorly understood. One experimentally characterized feature of STP is that it can be flexibly modulated by neuromodulators such as dopamine (Chiu et al., 2010; Gao et al., 2001; Gao et al., 2003; Kroener et al., 2009; Leyrer-Jackson and Thomas, 2018; Seamans et al., 2001a; Seamans et al., 2001b; Tecuapetla et al., 2007; Tritsch and Sabatini, 2012). Specifically, neuromodulators can alter the temporal profile of STP by governing release probability: enhancing initial release will more rapidly exhaust neurotransmitter vesicles from the readily releasable pool and favor short-term depression, in contrast, decreasing release probability can decrease short-term depression and favor short-term facilitation.

Even though STP is universally present at cortical synapses, most neural network models do not incorporate STP (for exceptions see (Buonomano and Merzenich, 1995; Masse et al., 2019; Mongillo et al., 2008; Murray and Escola, 2017)). And to the best of our knowledge, no previous neural network models have examined the computational role of the neuromodulation of STP. Here we demonstrate that the incorporation of STP, and its neuromodulation, into RNN models provides a powerful and flexible mechanism to temporally and spatially modulate RNN dynamics and thus sensorimotor control. We show that neuromodulation of STP accounts for experimental results on scaling tasks (Remington et al., 2018; Soares et al., 2016), and establish that while conventional RNNs can learn to temporally and spatially scale their dynamics, the incorporation of STP significantly enhances the ability of networks to generalize across temporal and spatial scales. Our results provide a novel hypothesis as to why synapses may exhibit STP, and provide a novel computational mechanism for unified spatial and temporal control of sensorimotor behavior.

## RESULTS

Firing-rate-based RNN models have successfully been used to capture neural dynamics of cortical circuits and account for how biological neural networks can perform a range of complex cognitive tasks (Chaisangmongkon et al., 2017; Laje and Buonomano, 2013; Mante et al., 2013; Murray et al., 2017; Sussillo and Abbott, 2009). With a few exceptions (Barak and Tsodyks, 2014; Masse et al., 2019), these RNN models generally do not incorporate STP. Here we incorporate STP in all the synapses of the RNN using the standard implementation composed of three variables(Markram et al., 1998): U, which can be interpreted as initial release probability or proportion of vesicles released from the readily-releasable-pool; τ_x_, the time constant of recovery from depression; τ_f_, the time constant of facilitation (**Fig. 1a**). We further implemented neuromodulation of STP via a factor α that modulated U. This factor α captures the level of a neuromodulator such as dopamine at the beginning of a trial. By modulating the U in STP through α, the STP can switch from short-term depression with high U values to short-term facilitation with low U values (**Fig. 1b**).

**Fig. 1:**
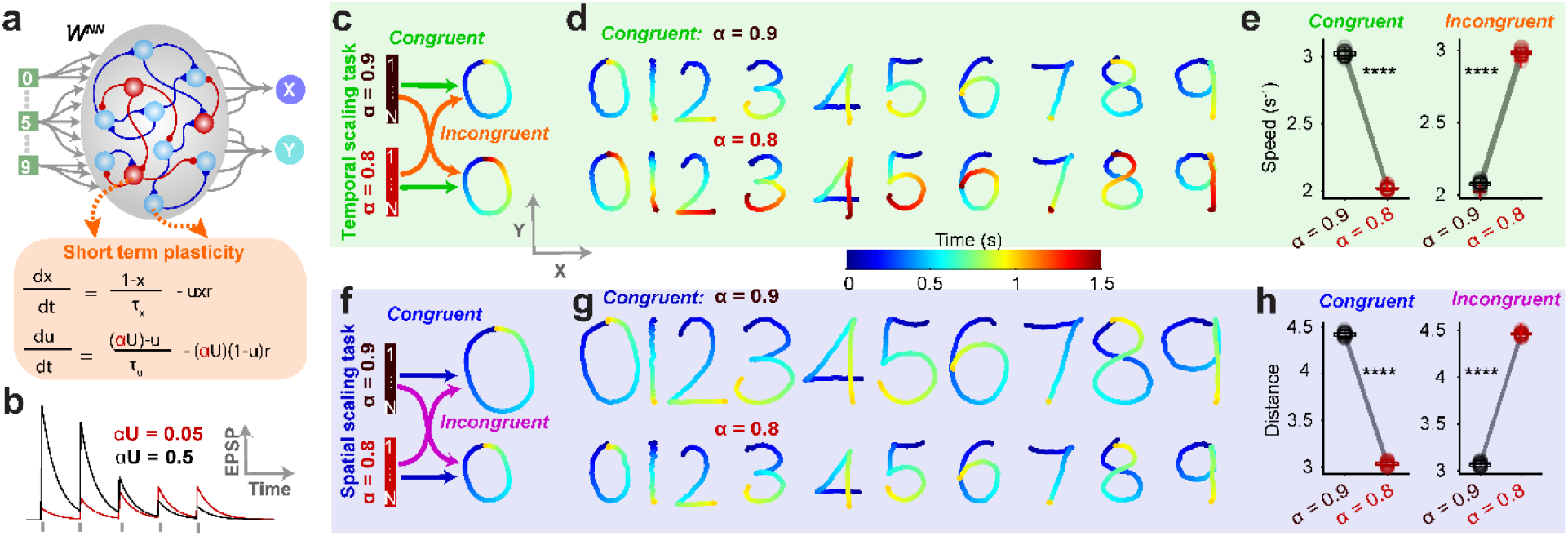
Temporal and spatial control of motor trajectories through neuromodulation of STP. **a** Schematic of the RNN with 80% excitatory (cyan) and 20% inhibitory (red) units. Transient activation of either of 10 inputs triggers the production of a digit (from 0 to 9). STP was implemented through the depression variable (x) and facilitation variable (u). For each trial, the constant U was scaled by α to signal temporal or spatial scale. **b**, Example of neuromodulation of STP. In the extreme, different values of αU can result in dramatic short-term depression or facilitation. A train of 20 Hz stimuli is delivered to the synapse, and τ_f_ and τ_d_ are both 1 s. **c**, Schematic of the temporal scaling task. For each trial, α was either 0.9 or 0.8 for all units, corresponding to a given digit production with a duration of 1 s and 1.5 s, respectively in the congruent condition, and 1.5 s and 1 s in the incongruent condition. The size of the target was the same for two α levels. **d**, Example output traces for all 10 digits under α = 0.9 (top) and α = 0.8 (bottom) for the congruent temporal scaling condition. **e**, Summary of the output speed (proportional to duration) averaged across digits under different α levels for the congruent (left) and incongruent (right) conditions for the temporal scaling task. Note that the speed under α = 0.9 case is around 1.5 times that of 0.8 in the congruent condition, and vice versa for the incongruent condition (n = 20 RNNs; P< 0.0001 two-sided Wilcoxon signed-rank test). **f**, Schematic of the spatial scaling task. For each trial, α was either 0.9 or 0.8 for all units corresponding to a scale of 1.5 and 1, respectively, in the congruent condition and 1 and 1.5, respectively, in the incongruent condition. The duration of the target was always 1 s. **g**, Example output traces for all 10 digits under α = 0.9 (top) and α = 0.8 (bottom) for the congruent spatial scaling task. **h**, Summary of the output distance traveled averaged across digits under different α levels for the congruent (left) and incongruent (right) conditions for the spatial scaling task. Note that the distance under α = 0.9 is around 1.5 times that under 0.8 for the congruent condition, and vice versa for the incongruent condition (n = 20 RNNs; P< 0.0001 two-sided Wilcoxon signed-rank test). Boxplot: central lines, median; bottom and top edges, lower and upper quartiles; whiskers, extremes; red cross, outliers.

Each RNN was trained to produce ten complex motor trajectories—handwritten digits 0-9—in response to one of ten brief inputs. In the temporal scaling task (**Fig. 1c**), RNNs were trained to produce each digit at a fast or slow speed, corresponding to a total duration of 1 or 1.5 s, respectively. The different speeds were cued by the values of α. In principle, α can control the speed of the output by higher values (0.9) cueing faster speeds and lower values (0.8) slower speeds (which we will refer to as the congruent condition); or conversely by higher and lower values cueing slower and faster speeds, respectively (incongruent condition). In both the congruent (**Fig. 1d**) and incongruent (**Extended Data Fig. 1a**) conditions, RNNs can learn the temporal scaling task equally well, as quantified by the speeds of trained output trajectories, which is 1.5 times faster at α = 0.9 than α = 0.8 in the congruent condition and vice versa for the incongruent condition (**Fig. 1e**). Training to criterion on the temporal scaling task was successful across a diverse range of hyperparameters including the mean time constants of the depression and facilitation, the pairing of α levels and scaling factors (**Extended Data Fig. 2a-c**).

For the spatial scaling task RNNs were trained to generate digits with the same duration, but with different spatial scales (1x and 1.5x). As in the temporal scaling task, the relationship between α and the scaling factor could be congruent (0.9/0.8 →1.5x/1x) or incongruent (0.9/0.8 →1x/1.5x) (**Fig. 1f**). In both the congruent (**Fig. 1g**) and incongruent (**Extended Data Fig. 1b**) conditions RNNs can learn the spatial scaling task well, as quantified by the Euclidian distance traveled by the output trajectories, which is 1.5 times more at α = 0.9 than α = 0.8 in the congruent condition and vice versa for the incongruent condition (**Fig. 1h**). Again, the training for the spatial scaling task was robust across a diverse range of hyperparameters (**Extended Data Fig. 2d-f**).

Although RNNs can learn equally well in both the congruent and incongruent conditions, the number of epochs needed to reach the same criterion for the congruent was significantly lower than that for the incongruent conditions in both the temporal and spatial scaling tasks (**Extended Data Fig. 1c)**. This implies that the congruent condition may offer intrinsic computational advantages (see below).

These results demonstrate that in principle, the α levels, which modulate STP through the initial “release ratio” can control either temporal or spatial scales, under both congruent and incongruent conditions. These results, however, do not address the more important question of how temporal and spatial scaling generalizes to novel values of α.

### Congruent modulation of STP generalizes better to novel scales in both temporal and spatial scaling tasks

To test how well temporal and spatial scaling generalizes to novel values of α, we tested the RNN performance under interpolated (α=0.8-0.9) and extrapolated (α<0.8, α>0.9) conditions by varying α values uniformly from 0.95 to 0.75 in both tasks. Optimal generalization would consist of output patterns that scaled linearly in time/space with α. For instance, in the congruent temporal scaling task, α = 0.85 should produce an output duration of 1.25 s, and α = 0.95 an output duration of 0.75 s. We quantified generalization as the RMSE of the actual outputs and the linearly scaled optimal targets. For the temporal scaling task RNNs generalized much better under the congruent compared to the incongruent condition (**Fig. 2a,c**). Additionally, the speed of the output trajectory scaled in a much more linear factor in the congruent condition (**Fig. 2d**). In the spatial scaling task, the generalization was also significantly better (but not as dramatically so) in the congruent condition (**Fig. 2b,e-f**).

**Fig. 2:**
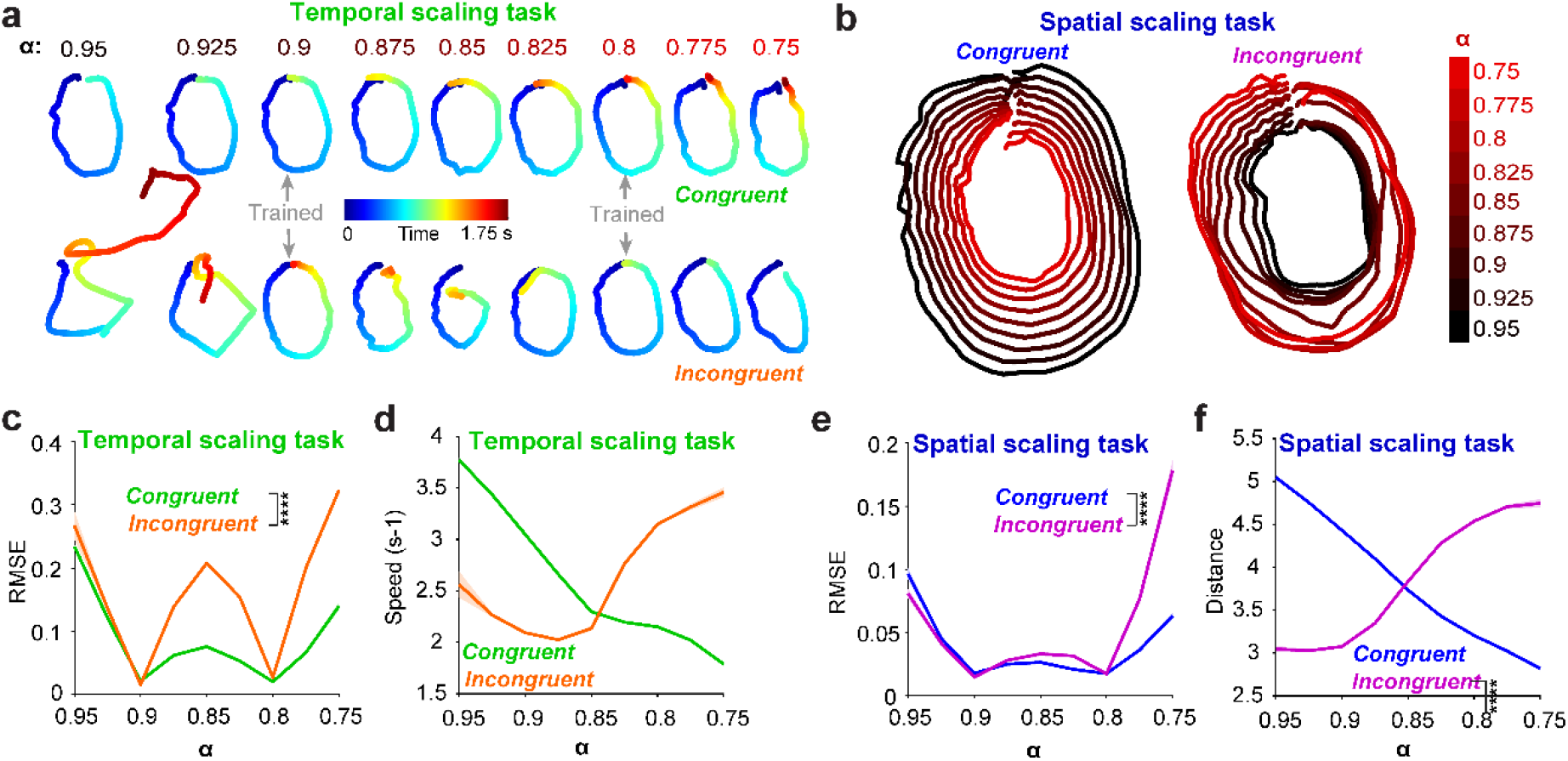
Generalization to novel scales is better in the congruent condition in both the temporal and spatial scaling tasks. **a**, Example output traces of digit 0 under novel α levels for congruent (top) and incongruent (bottom) conditions in the temporal scaling task. Gray arrows denote the α level used for training. **b**, Similar to **a** but for the spatial scaling task with congruent (left) and incongruent (right) conditions. Color codes different α levels. **c**, Summary of the generalization performance for the temporal scaling task as measured by RMSE between the actual output and targets linearly warped according to the corresponding α level. Note that RMSE at the novel (untrained) α values for the congruent (green) condition is significantly lower than that for the incongruent (orange) condition (n = 20 RNNs; two-way ANOVA with mixed-effect design, F_1,38_ = 707.7, P < 10^−25^). **d**, Summary of speed versus α levels for the congruent (green) and incongruent (orange) conditions in the temporal scaling task. Note that the relation for the congruent condition was more linear. **e**, Same as **c** but for the spatial scaling task. RMSE for the congruent (blue) condition was significantly lower than that for the incongruent (purple) condition (n = 20 RNNs; two-way ANOVA with mixed-effect design, F_1,38_ = 52.0, P < 10^−7^). **f**, Same as **d** but for distances with the congruent (blue) and incongruent (purple) conditions in the spatial scaling task. Data were presented as mean ± SEM (light overlay).

Although in both the congruent and incongruent conditions RNNs can be trained to the two target scales equally well, the generalization results indicate that the congruent relationship may be inherently better at modulating RNN dynamics and thus temporal and spatial scaling of output patterns.

### Temporal and spatial profile of recurrent dynamics

To begin to understand how neuromodulation of STP drives the scaling of RNN dynamics across temporal and spatial scales, we first analyzed the dynamics of the recurrent network under different values of α. We plotted the normalized population activity sorted by latency at different α values for congruent and incongruent conditions. For the temporal scaling task, the sequential order of the dynamics at α = 0.9 in the congruent condition preserved to dynamics at α = 0.8 but shifted to the left (**Fig. 3a left**, emphasized by the red line at 0.5 s). Visually, in the incongruent condition, there was a potential distortion of the sequence between α values of 0.9 and 0.8 (**Fig. 3a right**). This potential distortion can also be seen by plotting and comparing the two congruent RNN trajectories (non-normalized) and two incongruent trajectories in PCA space: the α = 0.9/0.8 trajectories appear to be more parallel in the congruent condition but not so in the incongruent condition (**Fig. 3b**). Visual inspection of the congruent and incongruent sorted neurograms in the spatial scaling task (**Fig. 3c**) are not as distinct as the temporal scaling task, in the sense that sequential order of the dynamics preserved across two α levels in both congruent and incongruent conditions. However, the non-normalized RNN trajectories in PCA space (**Fig. 3d**) revealed the distance traversed by the dynamics at α = 0.9 is larger than at α = 0.8. In other words, the size of the recurrent dynamics is ‘spatially larger’ at α = 0.9.

**Fig. 3:**
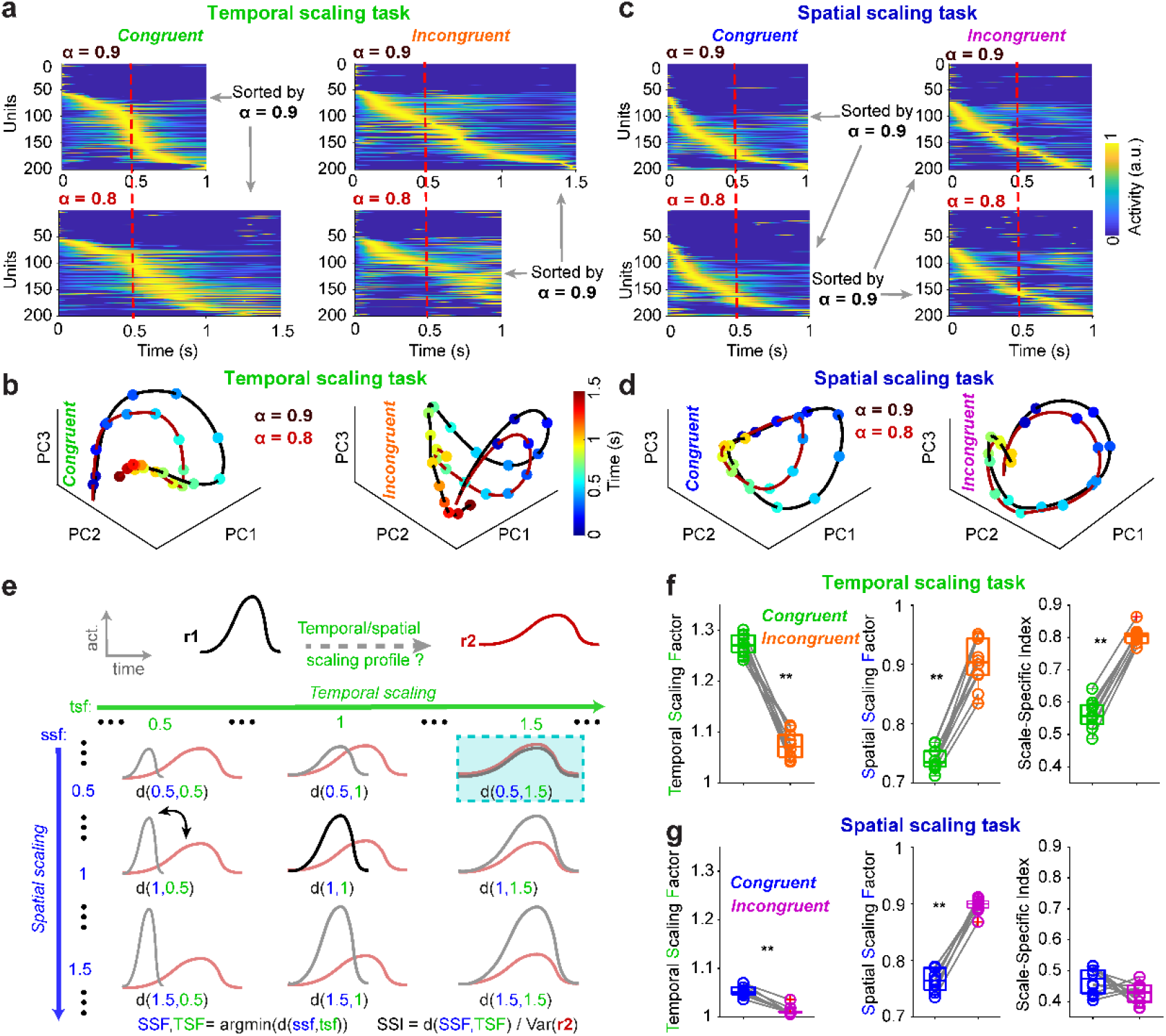
Temporal and spatial scaling of recurrent dynamics. **a**, Normalized recurrent population activity at α = 0.9 (top) and α = 0.8 (bottom) sorted according to the peak activity latency at α = 0.9 for congruent (left) and incongruent (right) temporal scaling task. For reference, the red dashed line denotes 0.5 s. **b**, Same as **a** but for spatial scaling task. **c**, Plot of the first three principal components of population activity at α = 0.9 (black) and α = 0.8 (dark red) for congruent (left) and incongruent (right) conditions in temporal scaling task. The color bar shows the code for time. **d**, Same as **c** but for spatial scaling task. **e**, Schematic of the calculation of the Temporal Scaling Factor (TSF), Spatial Scaling Factor (SSF), and Scale-Specific Index (SSI). For two hypothetic neural trajectories (illustrated as one dimension here), **r1** (black) and **r2** (red), the goal of the algorithm is to find the best temporal and spatial (amplitude) scaling factors by which warping **r1** provides the best match to **r2. r1** is temporal warped by either linear interpolating or subsampling a range of candidate temporal scaling factors (tsf). For each time-warped dynamics, we further multiply it with candidate spatial scaling (amplitude) factors (ssf), resulting in a grid of temporal-spatial warped dynamics of **r1** (gray). Finally, the average Euclidian distance between these warped dynamics and **r2** (light red traces on top) is computed. Thus, the distance is a function of tsf and ssf. The tsf and ssf leading to the minimal distance are defined as TSF and SSF respectively. The performance of the temporal-spatial profile was quantified by the SSI, which is the minimal distance at TSF/SSF divided by the distance between **r2** and its mean (similar to the variance of **r2**). **f**, comparison of congruent and incongruent conditions for average TSF (left), SSF (middle) and SSI (right) across 20 RNNs in the temporal scaling task (n = 10 digits; P = 0.002, two-sided Wilcoxon signed rank test for TSF, SSF, and SSI). **g**, Same as **f** but for spatial scaling task (n = 10 digits; P = 0.002, 0.002 and 0.131, two-sided Wilcoxon signed rank test for TSF, SSF and SSI respectively). Boxplot: central lines, median; bottom and top edges, lower and upper quartiles; whiskers, extremes; red cross, outliers.

To quantify the unified temporal and spatial (amplitude) scaling of the RNN trajectories, we developed three interrelated measures, Temporal Scaling Factor (TSF), Spatial Scaling Factor (SSF), and scale-specific index (SSI), all calculated from the same algorithm (schematized in **Fig. 3e**). The algorithm searches for the best temporal (TSF) and spatial (SSF) warping factors of a template RNN dynamics (e.g., α = 0.9; exemplified by the one-dimensional trace r1) that best matches the comparison trajectory (e.g., α = 0.8; exemplified by trace r2). The Euclidean distance between r1 and r2 at the best warping of r1 (at TSF and SSF) is then normalized (see Methods) to obtain an SSI value. Intuitively, the lower the SSI, the better r2 can be fitted through warping r1 temporally by TSF and spatially by SSF. Applying these measures to the temporal scaling task, we found that the TSF for the congruent condition is significantly higher than the incongruent condition (**Fig. 3f**, left). The SSI was also significantly lower (**Fig. 3f, right**), indicating that compared to the incongruent RNN trajectories, the congruent RNN trajectories at α=0.8 was a linearly warped version of the α=0.9 trajectory. Note that the SSF for the temporal scaling task is below 1 (**Fig. 3d, middle**), indicating that the faster trajectory is accompanied by higher amplitude firing rates of the RNN units.

Quantification of RNN trajectories in the spatial scaling task reveals that the SSF for the congruent condition is significantly lower than that for the incongruent conditions (**Fig. 3g, middle**) and that as expected the TSF for the spatial scaling task is close to 1 (**Fig. 3g**, left), which is consistent with the visual inspection in **Fig. 3d**. The low values of the SSI for both the congruent and incongruent trajectories indicate that in both conditions the α=0.8 trajectories are linearly warped versions of the α = 0.9 RNN trajectories.

To further corroborate the above measures we performed time and unit shuffled controls (**Extended Data Fig. 3**), in which shuffling resulted in significantly higher SSI suggesting an independent relationship between the two shuffled dynamics. We performed the same analyses based on the temporal profile of the synaptic efficacy as defined by the product of x and u in the STP model. This revealed similar temporal and spatial scaling profiles as the dynamics of activity (**Extended Data Fig. 4**).

In sum, in the temporal scaling task, the congruent neuromodulation of STP produced temporal scaling of output trajectories, by temporally scaling RNN trajectories. Importantly, for temporal scaling, there is a clear asymmetry between creating parallel RNN trajectories that unfold at slower speeds by either decreasing initial release strength (congruent) or increasing initial release strength (incongruent). In the spatial scaling task this asymmetry—i.e., the superior performance of decreasing initial synaptic strength to generate smaller output trajectories is less pronounced but still present.

### Mechanisms underlying the scaling of recurrent dynamics and output patterns

To dissect the mechanisms underlying the differential scaling of RNN trajectories by congruent and incongruent neuromodulation of STP, we focused on the RNN state trajectories at α = 0.9 and α = 0.8 (**s1** and **s2**, respectively, in **Fig. 4a)**. At any given time point in **s1** and **s2** with **p12** being the direction from **s2** to **s1**, there is a velocity vector **v1** and **v2**, respectively, that can be decomposed into the decay component (**d1** and **d2**) and a recurrent component (**rec1** and **rec2**). We defined the angle between **rec2** and **p12** as *θ* and the angle between **rec2** and **v1** as *μ*. Intuitively, for **s1** to speed up compared to **s2**, as in the congruent temporal scaling task, we would expect the **rec2** and **v1** to point in a similar direction—with the angle between the two (*μ*) smaller than 90 degrees—to increase the drive on the **v1** direction, and vice versa for the incongruent condition. On the other hand, for **s1** to transition to a larger trajectory, we would expect the **rec2** and **p12** to point in a similar direction—with the angle between the two (*θ*) to be smaller than 90 degrees—in order to increase drive on the **p12** direction.

**Fig. 4:**
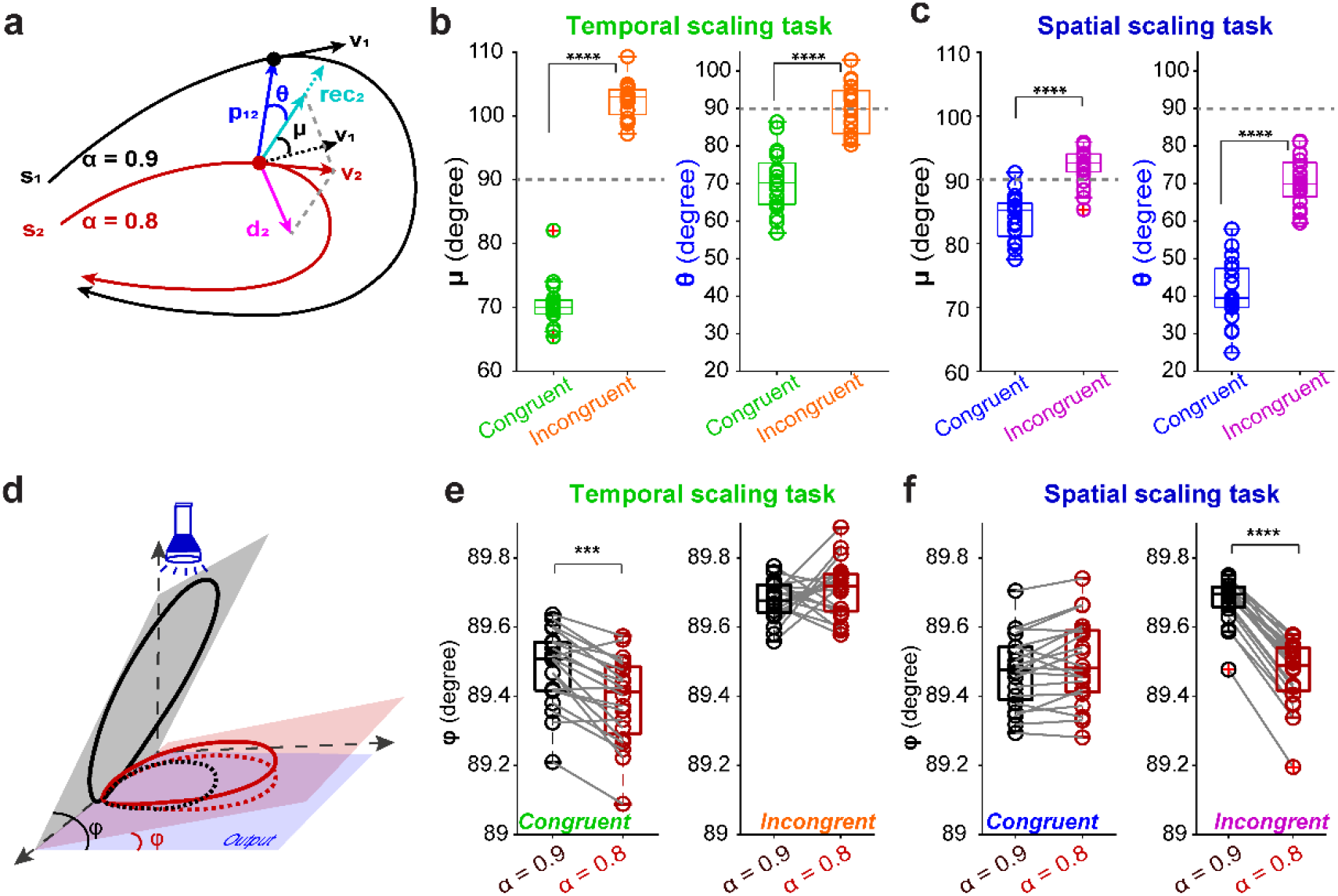
Subspace analysis of the temporal and spatial scaling of RNN dynamics and output. **a**, Schematic of the decomposition analysis between recurrent activity states at α = 0.8 (s2, red curve) and α = 0.9 (s1, black). For a given time point on s2 (red dot) and its corresponding point (normalized time) on s1 (black dot) there are velocity vectors v2 (red arrow), and v1 (black arrow), respectively. v2 can be decomposed into the recurrent component, rec2 (cyan arrow), and decay component d2 (magenta arrow). θ denotes the angle between rec2 and p12 and μ denotes the angle between rec2 and v1. **b**, The average angle μ (left) and θ across time in the temporal scaling task. μ is significantly lower for the congruent settings than for the incongruent setting (n = 20 RNNs; P < 10^−7^, two-sided Wilcoxon rank sum test), while θ is slightly, but significantly lower for the congruent settings than for the incongruent task as well (n = 20 RNNs; P < 10^−6^, two-sided Wilcoxon rank sum test) **c**, same as **b** but for spatial scaling task. θ is significantly lower for the congruent settings than that for the incongruent setting (n = 20 RNNs; P < 10^−7^, two-sided Wilcoxon rank sum test). while μ is slightly but significantly lower for the congruent settings than for the incongruent setting as well (n = 20 RNNs; P < 10^−6^, two-sided Wilcoxon rank sum test). **d**, Schematic of how recurrent dynamics with smaller (red solid) or larger (black solid) size can generate larger (red dashed) or smaller (black dashed) outputs in output space, respectively. **e**, Comparison of the angle between the planes of recurrent space (2 PCs) and output space at different α levels for congruent (left) and incongruent(right) cases in the temporal scaling task (n = 20 RNNs; P< 0.001 and P = 0.263 for congruent and incongruent respectively, two-sided Wilcoxon signed-rank test). **f**, Same as **e** but for the spatial scaling task (n = 20 RNNs; P= 0.062 and P < 0.0001 for congruent and incongruent respectively, two-sided Wilcoxon signed-rank test). Boxplot: central lines, median; bottom and top edges, lower and upper quartiles; whiskers, extremes; red cross, outliers.

Indeed, for the temporal scaling task, the mean *μ* across time in the congruent condition is lower than 90 degrees, significantly lower than in the incongruent condition that is larger than 90 degrees (**Fig. 4b, left)**. *θ* in the congruent condition for the temporal scaling task is significantly lower than in the incongruent condition (**Fig. 4b, right**), which is consistent with the SSF in congruent conditions being slightly lower than in incongruent conditions (**Fig. 3g, middle**). In the spatial scaling task, *θ* in the congruent condition is significantly lower than in the incongruent condition (**Fig. 4c, right**), which is consistent with the results that SSF in the congruent spatial scaling task is significantly lower than in the incongruent conditions (**Fig. 3g, middle**). While μ in both congruent and incongruent spatial scaling tasks are close to 90 degrees, but with congruent conditions being slightly slower, which is also consistent with the TSF in congruent conditions being slightly higher (**Fig. 3g, left**).

As shown in **Fig. 3**, SSFs of the recurrent dynamics in both the congruent and incongruent conditions are less than 1, which indicates that the size of the recurrent dynamics at α = 0.9 is larger than α = 0.8. The larger recurrent dynamics at α = 0.9 in the congruent spatial scaling task would be appropriate for generating larger output as the task requires. However, in the incongruent condition, this scenario would change so that a larger recurrent dynamics needs to generate a smaller output. This paradox exists for the temporal scaling task too, where the larger recurrent dynamics is meant to generate an output of the same size as the smaller one. The resolution to this apparent paradox can be understood using a light projection analogy (**Fig. 4d**). In light projection, to get a bigger shadow from a smaller trajectory, one can arrange the smaller trajectory within a smaller angle with the ground, the plane of the shadow. Analogously, the motor output is the projection of the recurrent dynamics onto the output space governed by the output weights. Thus, we would expect the angle between smaller recurrent dynamics (subspace of the first 2 PCs) and output space to be smaller than that between larger recurrent dynamics and output space. Indeed, the angle between recurrent space at α = 0.8 and the output space was slightly but significantly lower than that at α = 0.9 in the congruent temporal scaling task (**Fig. 4e left**) and incongruent spatial scaling task (**Fig. 4f right**). These differences were not as significant in the incongruent temporal scaling task probably due to the higher SSF (**Fig. 4e right**), or in the congruent spatial scaling task (**Fig. 4f left**) as one would expect (no space alignment required for pairing bigger recurrent dynamics with bigger output). Those findings are robust when quantifying the angles for higher dimensional recurrent space expanded by more PCs (**Extended Data Fig. 4**).

### Short-term plasticity enhances generalization and speeds up training

The above results demonstrate that we can modulate the temporal or spatial scale by changing the initial synaptic release probability controlled by α. Although α controlled release probability U we didn’t directly address whether STP is actually contributing to the results. That is, does simply adjusting the synaptic release probability in the absence of STP (i.e., simply scaling all synaptic weights) result in similar performance? To address that question, we ran control simulations for RNNs without STP but still included a α term. Specifically, we fixed *x* at 1 and *u* at αU during the whole trial. These modifications removed the STP dynamics but keep the overall magnitude of the connection weights at the same scale as the control. We then trained and tested the RNNs without STP doing the same task in the congruent condition as the standard model. Interestingly, the generalization performance for the RNNs without STP dramatically decreased in both the temporal (**Fig. 5a,c**) and spatial scaling tasks (**Fig. 5b,c**) compared to the RNNs with STP. Furthermore, STP dramatically speeded up training as shown by fewer training epochs needed to reach the same criterion (**Fig. 5d)**. Thes results suggest that STP does indeed provide a novel mechanism to effectively scale temporal and spatial neural dynamics.

**Fig. 5:**
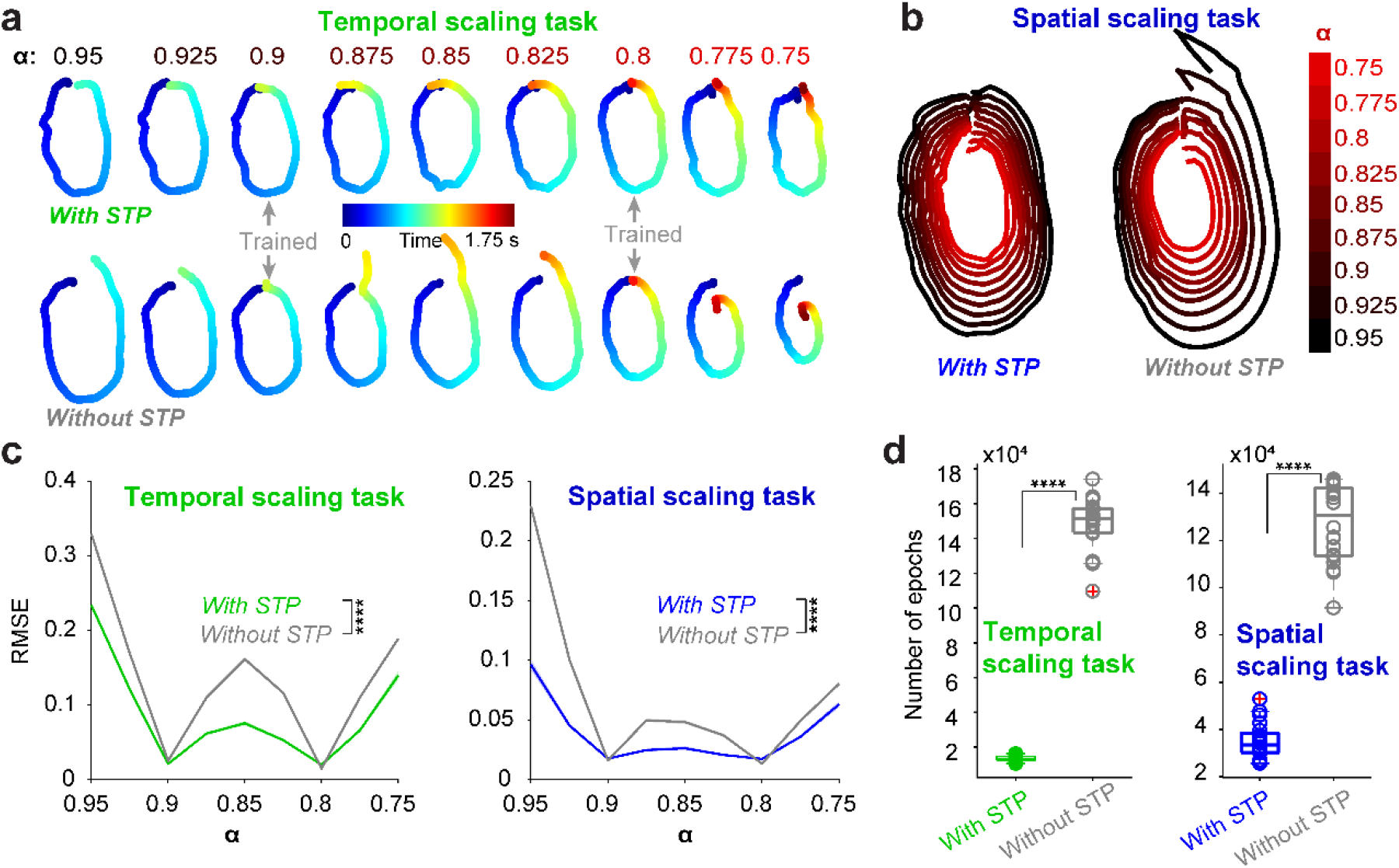
STP enhances generalization performance and speeds-up learning. **a**, Example output traces of digit 0 under different α levels for congruent temporal scaling task with STP (top) and congruent condition without STP (bottom). Gray arrows denote the α level used for training. **b**, Same as **a** but for the spatial scaling task. Color codes for different α levels. **c**, Summary of the generalization performance by the average RMSE between output and the linearly scaled targets. Note that in both temporal (left) and spatial (right) scaling tasks the RMSE for the novel α levels for the RNNs with STP (green or blue) case was significantly lower than that for RNNs without STP (gray) (n = 20 RNNs; two-way ANOVA with mixed-effect design, F_1,38_ = 1102.4, P < 10^−28^ and F_1,38_ = 556.3, P < 10^−23^ for temporal and spatial scaling task respectively). **d**, Comparison of the number of training epochs to reach criteria for RNNs with STP and without STP in temporal scaling task (left) and spatial scaling task right (n = 20 RNNs; P < 10^−7^, two-sided Wilcoxon rank sum test for both tasks). Boxplot: central lines, median; bottom and top edges, lower and upper quartiles; whiskers, extremes; red cross, outliers.

### Joint control of temporal and spatial scales, and shape via neuromodulation of STP

Up to now, we have demonstrated that adjusting α can control either temporal or spatial scales separately. We next ask whether α jointly controls the temporal and spatial scale in a single RNN. To explore this possibility, we arbitrarily divided the recurrent units into two groups. One group receives a neuromodulatory signal that will control temporal scale by altering α, and another group receives a signal that controls spatial scale (**Fig. 6a**). RNNs can learn this task well as shown (**Fig. 6b**). And importantly, in this joint control task, RNNs can generalize well to all the combinations of novel α levels for the temporal and spatial scales supported by the speed and distance of the outputs (**Fig. 6c**). Note that speed progressively increases towards to the top left part of the speed plot, which corresponds to the fastest speed required to generate the largest output in the shortest time. Similarly, the distance progressively increases towards the bottom left part of the distance plot, which corresponds to the longest distance required to generate the largest output in the longest time.

**Fig. 6:**
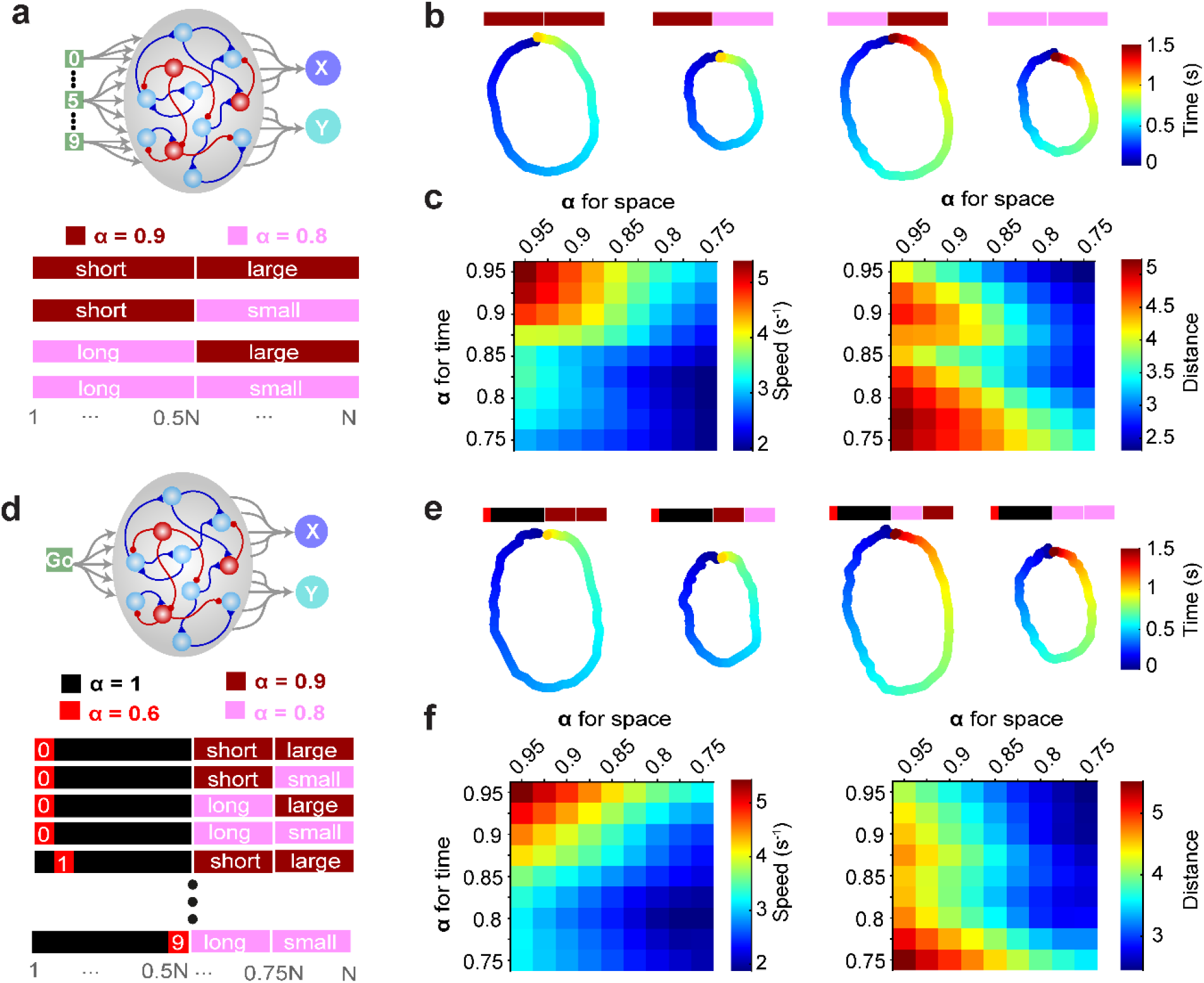
Joint control of temporal and spatial scales in RNNs through differential modulation of α in distinct subpopulations. **a**, Schematic of unified control of temporal and spatial scale. The α level of 50% selected units controls temporal scaling, while the other half spatial scaling. **b**, Example output traces for digit 0 in four cases: short-large, short-small, long-large, and long-small. **c**, Summary of average speed (left) and distance (right) across different α levels. **d**, Schematic of encoding temporal and spatial scale, and digit identity through α. 50% units to signal digits identity, while half of the rest 50% units signal duration (25%) and spatial scale (25%). For a given digit, 20 randomly selected units (out of 400) were assigned α = 0.6 among the digit population, while α = 0.9/0.8 were used for the temporal and spatial scaling as in the above model. **e**, Same as **b** but for the model in **d. f**, Same as **c** but for the model in **d**.

We next asked whether α can jointly control an additional task dimension: the output shape, i.e., the identity of the digit was cued by changing α in a subset of units rather than distinct inputs. To achieve this, we divided the recurrent units into three groups (50%, 25%, 25%): 50% for digit shape, 25% for temporal scale, and the rest 25% for spatial scale (**Fig. 6d)**. The digit group was further divided into ten subgroups with the α of each set to 0.6 and the rest 1 to signal each of the ten digits (**Fig. 6d)**. With this architecture, RNNs learned and generalize well to joint control of temporal and spatial scales while also cueing different digits with α (**Fig. 6e,f**).

Although the two strategies for joint control—using either different inputs or α signaling for digit identity—exhibited similar learning and generalization performance, PCA plots revealed that using α signaling the digit seemed to lead to recurrent dynamics more similar across digits (**Extended Data Fig. 6a**). These visual results are further confirmed by the cross-digit correlations for the two strategies (**Extended Data Fig. 6b, c**). These findings suggest that similar output can be generated from different recurrent dynamical regimes, similar to what we observed in the congruent vs incongruent conditions.

### Neuromodulation of STP captures the temporal scale observed in two sensorimotor tasks

For the analysis above, we mainly focused on how α can adjust the temporal and spatial scale in a motor control task—generating digit handwriting. To investigate whether modulating α can account for experimental funding on temporal scaling we simulated two experimental studies: one from rodents and another from non-human primates. First, we trained RNNs to solve an interval-alternative-forced-choice task (IAFC) where rats needed to classify intervals as short or long (Soares et al., 2016). In this study optogenetically increasing dopamine levels selectively shifted the decision towards the short intervals (i.e., the psychophysics of long choice probability was shifted right). To replicate this experiment, we first trained RNNs to do the same interval discrimination task with a single α = 0.8 for all units (**Fig. 7a**). The RNNs can replicate the behavioral results as shown by the output traces for an example RNN (**Fig. 7b**) and the psychophysics curve of the long choice probability (**Fig. 7c)**. We then sought to simulate the dopamine manipulation experiments in our model. Multiple studies have demonstrated that dopamine decreases the synaptic release probability in both excitatory and inhibitory cortical synapses (Chiu et al., 2010; Gao et al., 2001; Gao et al., 2003; Seamans et al., 2001b; Tritsch and Sabatini, 2012), we thus simulated dopamine levels in the IAFC task at either α = 0.9 or 0.7 to capture low or high dopamine levels, respectively. Decreasing α from 0.8 to 0.7—which corresponds to increasing dopamine—shifts the psychophysics curve of the long choice probability to right, the same as the results of dopamine manipulation experiments, and vice versa for increasing α to 0.9 (**Fig. 7d,e)**. These results are consistent with the congruent temporal scaling motor trajectory task, in the sense that in both cases, decreasing α slows down the recurrent dynamics, and thus slows down either the motor output in the motor trajectory task or sensory timing in the IAFC task resulting underestimating the input intervals.

**Fig. 7:**
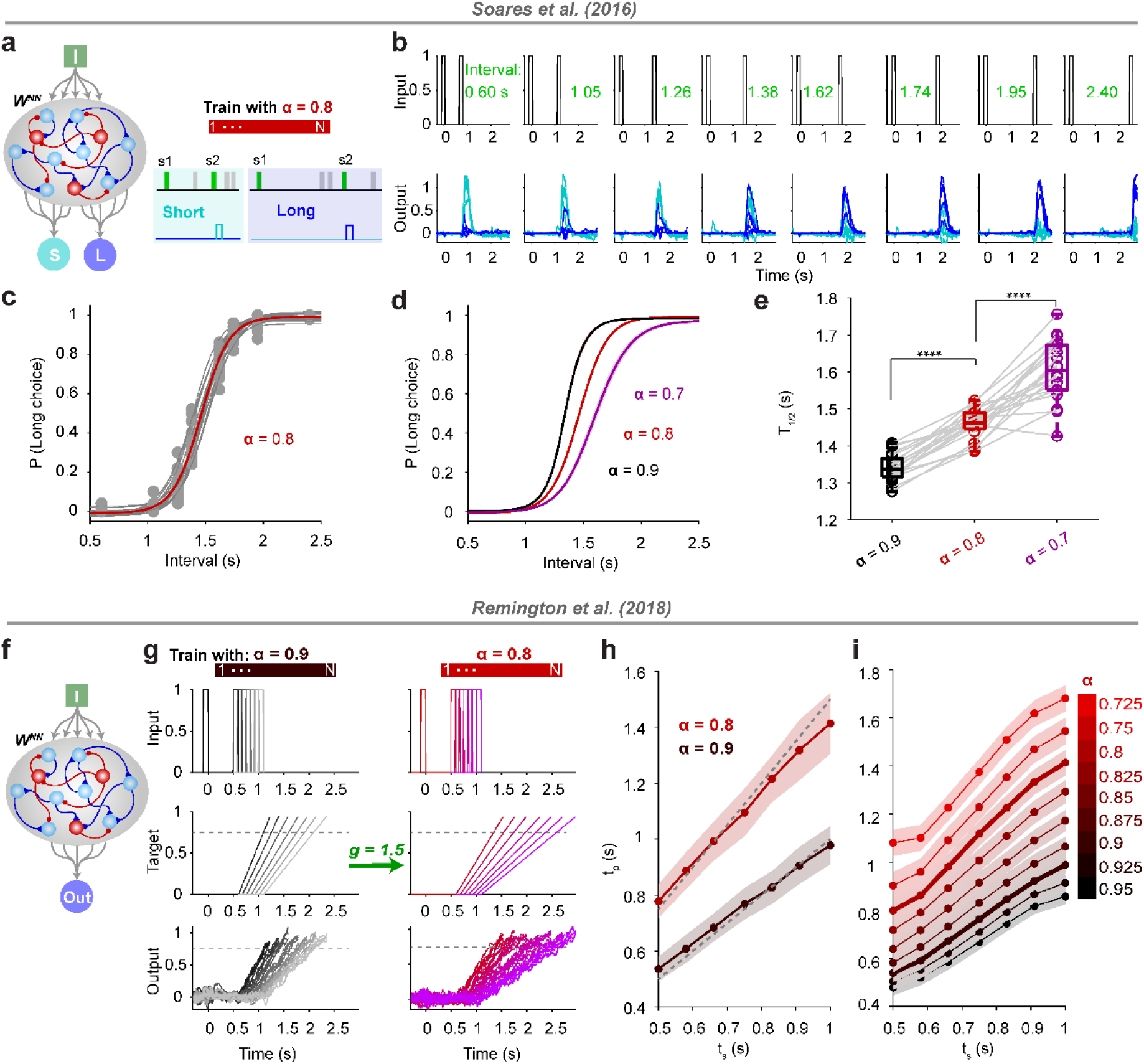
Neuromodulation of STP captures the experimental results of two sensorimotor timing tasks. **a**, Schematic of RNN used to simulate the interval alternative-forced choice task of Soares et al (2016). RNNs are composed of one input for delivering two stimuli with a range of intervals between 0.6 and 2.4 s (short < 1.5 s; long for > 1.5 s) and two outputs corresponding to a categorical short or long decision. RNN were trained only with α = 0.8 for all units. **b**, Output traces of an example RNN for all the intervals tested. **c**, Sigmoidal fits of the long choice probability for 20 RNNs tested at α = 0.8. **d**, Similar to **c**, but testing the network at α = 0.9 (black), α = 0.7 (purple) and α = 0.8 (red) for comparison. **e**, Summary of the time “point-of-subjective equality” (T_1/2_) for the sigmoid fits in **d**. Changing α significantly changed the T_1/2_ (n = 20 RNNs, Kruskal-Wallis test, P<10-10, χ^2^_(2,57)_ = 48.0) and T_1/2_ for α = 0.9 and 0.7 is significantly lower and higher than that for α = 0.8 respectively (P = 0.0007 and P = 003 respectively, by Dunn’s multiple comparison test). **f**, Schematic of RNN used to simulate a Ready-Set-Go task of Remington et al (2018). RNN was composed of one input that delivered to events (demarcating the Ready-Set interval) and one output. Based on the context, cued by α = 0.9 or α = 0.8, the output unit should generate a duration of 1x or 1.5x the Ready-Set interval, the production time. **g**, Plot of the input (top), target (middle) and output traces for α = 0.9 (left) and α = 0.8 for an example RNN. The Gray dashed line denoted the threshold used to quantify the crossing time. **h**, Plot of the production time vs the sensory time for α = 0.9 (black) or α = 0.8 (red) in one example RNN. Data were presented as mean ± SD. **i**, Summary of the average production time for novel α levels with trained α shown thicker lines (n = 20 RNNs). Data were presented as mean ± SEM.

We next simulate an experimental task that used the Ready-Set-Go paradigm in monkeys. In this task, subjects perceive an initial interval demarcated by ready and set cues, and they have to produce a go response in which the set-go interval is the same as the ready-set interval. In this experimental study (Remington et al., 2018) there was an additional scaling cue, which determined if subjects were required to produce the exact ready-set interval (1x) or scale the interval by 1.5x. To simulate this flexible sensorimotor timing task, we trained RNNs to do the task the same as the experiment with α = 0.9/0.8 signaling the 1x/1.5x context respectively (**Fig. 7f,g)**. RNNs learned this task well and captured some important features of the behavior, such as the regression to the mean effect—bias of the long/short interval towards the mean (**Fig. 7g,h**).

Similar to the motor trajectory task, RNNs generalize well to the novel α levels with the scaling factor changing smoothly from the trained ones, 1x/1.5x (**Fig. 7i**).

## DISCUSSION

Here we have proposed, and provided support, for the hypothesis that neuromodulation of STP provides a mechanism for recurrent neural networks to scale their dynamics in both time and space. RNNs that incorporated STP and used neuromodulation of STP to signal changes in temporal and/or spatial scale exhibited better generalization and performance than RNN models in which temporal and spatial scales were signaled by distinct inputs (**Extended Data Figure 8**) or changes in absolute synaptic weights alone (**Fig. 5**). Furthermore, neuromodulation of STP allowed RNNs to capture the results of two experimental studies based on distinct sensorimotor timing tasks. While neuromodulation of STP is a well-established experimental phenomenon in cortical and subcortical circuits alike (Baimoukhametova et al., 2004; Burke et al., 2018; Gonzalez-Islas and Hablitz, 2003; Leyrer-Jackson and Thomas, 2018; Nadim and Bucher, 2014; Rush et al., 2002; Seamans et al., 2001a; Tecuapetla et al., 2007), its potential role in neurocomputation has not been addressed. Here we establish that it provides novel mechanisms for the flexible regulation of neural dynamics and thus of motor control.

### Scaling of neural dynamics through neuromodulation of STP

Neuromodulators such as dopamine have been implicated in a large range of cognitive functions including reinforcement learning (Schultz et al., 1997) and timing. In the case of timing it has been proposed that DA may alter clock speed (Buhusi and Meck, 2002; Meck, 1996; Soares et al., 2016). How DA could alter the speed of the neural clock at the neural level, however, has not been addressed. Here we propose that DA’s ability to modulate STP provides a novel mechanism to link findings at the neural and cognitive levels. Specifically, in cortical circuits, synaptic transmission studies indicate that DA often, but not always, decreases EPSP amplitude through synaptic release probability (Burke et al., 2018; Leyrer-Jackson and Thomas, 2018; Seamans et al., 2001a; Tritsch and Sabatini, 2012). By incorporating STP into RNNs, and emulating dopaminergic inhibition of release probability through decreases in the variable α we were able to link cellular-level observations with previous systems and behavioral-level results (Soares et al., 2016).

We emphasize, however, that dopaminergic modulation of synaptic transmission is complex and dependent on brain areas and synapse classes (Burke et al., 2018; Leyrer-Jackson and Thomas, 2018; Nadim and Bucher, 2014). Similarly, at the behavioral and cognitive levels, the effects of dopamine are also highly complex. In the motor domain, for example, dopamine has been demonstrated to increase movement speed (amplitude) through modulation of the striatal activity(Panigrahi et al., 2015), potentially, reflecting differences in dopaminergic neuromodulation in different brain areas.

### Temporal versus spatial scaling

The ability of neuromodulation of STP to control temporal and spatial RNN dynamics was not the same. Our results suggest that neuromodulation of STP is better suited to control temporal, compared to spatial dynamics. Specifically, the difference between the congruent and incongruent conditions in the temporal scaling task was more distinct than in the spatial scaling task (**Fig 2**), as was the difference between the STP and control conditions in which scale was cued by synaptic strength in the absence of STP (**Fig. 5)**.

Previous studies have also shown that RNNs can account for temporal scale by cueing speed through the amplitude of a tonic “speed” input (Hardy et al., 2018; Remington et al., 2018; Saxena et al., 2022; Wang et al., 2018; Zhou et al., 2022) or by altering the gain though changes in intrinsic excitability (Lindén et al., 2022; Stroud et al., 2018)(an approach similar to our control condition in which all RNN weights were scaled). As in the current study, in these previous studies, temporal scaling was achieved by creating parallel neural trajectories that flowed at different speeds. We also directly compared previous approaches with neuromodulation of STP by implementing the input-cued mechanism in RNNs but in the absence of STP. As with the overall modulation of RNN weights control (**Fig. 5**), we found that the generalization and performance for the temporal scaling task were significantly worse than that for the α-cued mechanism (**Extended Data Fig. 8d**) but less different for the spatial scaling task (**Extended Data Fig. 8e**). These results were further confirmed by training RNNs at only a single speed level, which demonstrated that neuromodulation of STP exhibited better generalization to novel input levels (**Extended Data Fig. 8f**,**g)**.

### Dopamine and temporal scaling

The finding that neuromodulation of STP may serve as a neural mechanism for temporal scaling is consistent with the fact that DA has been linked to timing for decades (Fung et al., 2021; Maricq and Church, 1983; Meck, 1996; Rammsayer, 1999). However, there is a long-standing debate as to the direction of this relationship, which has remained a point of controversy (Simen and Matell, 2016). Specifically, early reports suggested that neuropharmacologically enhancing dopamine levels accelerated the neural clock (Buhusi and Meck, 2002; Lake and Meck, 2013), or rather suggested that there was no consistent effect or that DA accelerated the neural clock (Drew et al., 2003; Soares et al., 2016). Assuming that DA acts in part by decreasing synaptic strength and/or release probability—as in the canonical case of dopaminergic neuromodulation (Burke et al., 2018; Seamans et al., 2001a), our results strongly predict that DA should slow the internal clock. Specifically, as shown in **Fig 2**, the congruent relationship between decreasing the probability of release and slowing the clock is a significantly better way to temporally scale neural dynamics compared to the incongruent case where a neuromodulator such as DA would decrease clock speed.

Overall, our results demonstrated that the incorporation of STP in recurrent neural network models, and its neuromodulation, provides a powerful and flexible mechanism to implement temporal scaling as well as spatial scaling. These results thus provide a novel hypothesis as to why synapses may exhibit STP, and provide a novel computational mechanism for temporal and spatial scaling of neural dynamics, and thus of temporal and spatial control of sensorimotor tasks.

## METHODS

### Recurrent neural network model

#### Network architecture and STP

As in **Fig. 1**, RNNs were based on firing-rate units that obeyed Dale’s law (N = 200 unless otherwise specified, 80/20% excitatory/inhibitory). RNN dynamics was described by the following equations:

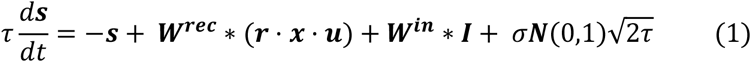

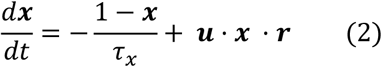

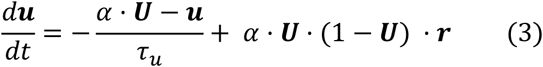

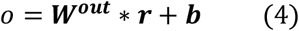

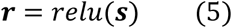

where **s** ∈ℝ^N×1^ represents the state of the RNN units, and the firing rate vector **r** corresponds to the rectified linear activation function on **s**. The time constant τ was 100 ms for all units. **W**^**in**^ ∈ℝ^N×2^ and **I** represent the input weights and external inputs. Each unit received independent Gaussian noise **N**(0,1) with the standard deviation of 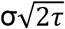. Unless otherwise specified, σ = 0.01. **W**^**rec**^ ∈ℝ^N×N^ is the recurrent weight matrix. Self-connections were absent in the network. Star represents the matrix product and the dot represents the element-wise product.

STP was incorporated as in previous models (Masse et al., 2019; Tsodyks and Markram, 1997). Cell-specific STP was implemented in the recurrent units as described in equations (2-3). Specifically, the depression variable **x** and facilitating variable *u* were shared for all synapses from the same pre-synaptic neuron. The vector **U** corresponds to the initial synaptic release probability or baseline percentage of available transmitter released. To implement neuromodulation of STP we scaled U with a factor α in the range of 0-1.

The output (o) of the network is computed linearly from the output weights Wout and r with a bias term b. RNNs were implemented and trained in Tensorflow 2.3 based on the code from a previous study (Kim et al., 2019).

#### Training

Networks were trained using adaptive moment estimation stochastic gradient descent algorithm (Adam) implemented in Tensorflow2 to minimize the RMSE (root mean square error) between network output o and target z:

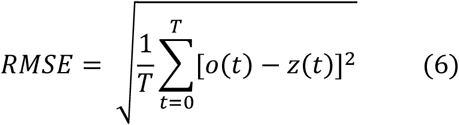

where T is the total length of a given trial. The target is task-dependent as described below. The learning rate was 0.001, and other TensorFlow default values were used. A discretization step of 10 ms was used for the simulations.

**W**^**rec**^ was initialized as a random matrix with full connectivity from a Gamma distribution with a shape parameter of 0.1 and scale parameter of 1.0, multiplied by a gain factor of 0.5. In order to start from an approximately balanced regime the inhibitory weights were multiplied by 4. To respect Dale’s law during training a rectified linear operation was applied on **W**^**rec**^ to clip the weights at zero and then excitation and inhibition were implemented by multiplying the clipped **W**^**rec**^ with a diagonal matrix of 1 and -1 representing excitatory and inhibitory units, respectively. Initial **W**^**in**^ was drawn from the same Gamma distribution clipped to zero during training the same way as **W**^**rec**^. **W**^**out**^ and **b** were initialized as zero.

**U** was drawn from a Gaussian distribution with a mean of 0.5 and standard deviation of 0.17 (mean/3) and cut off at 0.001 and 0.99. Unless otherwise specified τ_x_ and τ_u_ were drawn from a Gaussian distribution with a mean 1 s and standard deviation of 0.33 s (mean/3) and cutoff at 0.1 s and 3 s to ensure numerical stability. α was task-specific as described below.

Only **W**^**in**^, **W**^**rec**^, **W**^**out**,^ and **b** were trained. Parameters were updated after each batch of 16 trials. After every 100 batches of training, the network was tested for 20 batches to compute the task performance or mean error. For all the spatial and temporal scaling motor trajectory tasks, the training was considered a success and stopped when the mean error is lower than 0.02; while for the interval-alternative-forced-choice (IAFC) and flexible sensorimotor timing tasks (as described below), the criterion would be the task performance being higher than 90% or 98% respectively to capture the experimental features.

### Temporal and spatial scaling motor trajectory task

We trained RNNs to generate a series of complex motor trajectories: 10 handwritten digits from 0 to 9 (Goudar and Buonomano, 2018) of length 1 s. Unless otherwise specified, each of the ten inputs was presented for 0.1 s with onset time randomly drawn from a uniform distribution (0.2-s) to signal the digit identity. Following the input, the network evolved freely for a specific duration to match the corresponding targets warped temporally and/or spatially according to the task requirements.

#### Temporal scaling task

During training, for the congruent condition, α = 0.9 corresponded to the standard target with a duration of 1 s, while α = 0.8 trials corresponded to the target with a duration of 1.5 s uniformly interpolated from the standard one. For the incongruent condition, the association between the α and target duration switched, namely 0.9 and 0.8 corresponding to 1.5 and 1 s, respectively.

#### Spatial scaling task

During training, for the congruent condition, α = 0.8 corresponded to the standard size target with a duration of 1 s, while α = 0.9 corresponded to the larger target also with a duration of 1 s but with the amplitude multiplied by a factor of 1.5, namely the size of target was 1.5x larger than the standard one. For the incongruent condition, the association between the α and target size reversed.

#### Joint temporal-spatial scaling task

For the joint control of temporal and spatial scales in the same network (**Fig. 6**) we increased N to 400. 50% of recurrent units were used for temporal scaling control and the other 50% for spatial scale control (**Fig. 6a**).

To examine joint control the shape (digit identity), and temporal and spatial scale in a single network, we randomly divided the 400 recurrent units into three groups: shape (1-200), temporal scaling (201-300) and spatial scale (301-400). The temporal and spatial scale control was implemented the same way as the joint control network with inputs signaling the digit. We further divided the shape group into 10 subgroups corresponding to the 10 digits. For a given digit, we set the α in the corresponding subgroup to 0.6 while leaving the α of the rest of the shape group being 1 (**Fig. 6d**).

### Generalization performance

To see if RNNs trained with two α levels corresponding to two temporal or spatial scales can generalize to other scales, we test the RNNs trained with tasks of temporal, spatial scaling, or joint control of both with α level in between (interpolation) or outside (extrapolation) of the trained levels. Specifically, for trained level α1/α2 (for example, 0.9/0.8 for congruent conditions), we tested α = 0.95, 0.925, 0.875, 0.85, 0.825, 0.775 and 0.75. The generalization performance was quantified as the RMSE between the output under the testing α and the target z_α_(t) uniformly warped according to the corresponding trained α1/α2. For instance, the length of z_α_(t), T_α_ for the congruent temporal scaling task with the trained α1 and α2 values corresponding to the digit lengths T1 and T2 (1, 1.5 s) would be:

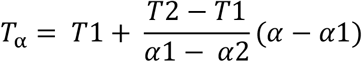

Therefore

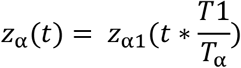

Similarly, the size of z_α_(t), S_α_ for the congruent spatial scaling task with the trained α1/α2 corresponding to the digit size S1/S2 (1.5/1) would be:

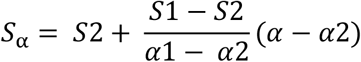

Therefore,

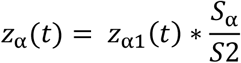

The incongruent conditions were modified accordingly.

### TSF, SSF, SSI

To quantify temporal and spatial scaling of the recurrent dynamics we extended a previously described method (Zhou et al., 2020, 2022) to define three measures: Temporal Scaling Factor (TSF), Spatial Scaling Factor (SSF), and the scaling-specific index (SSI). As in **Fig. 6d**, for two given population trajectories: **r1**(N×T1) and **r2** (N×T2) with T1<T2, the goal of the algorithm is to find the best temporal and spatial scaling factors, by which warping **r1** gives the best match to **r2**. Specifically, we searched among a range of temporal scaling factors (tsf, 0.5-2), and spatial scaling factors (ssf, 0.5-2). We then warped r1 temporally and spatially as follows:

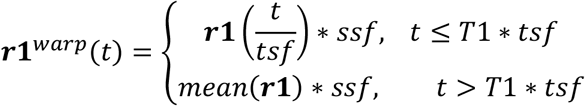

where the mean() function is applied to each unit.

To compare with **r1**^warp^(t), we extended the **r2** dynamics as follows:

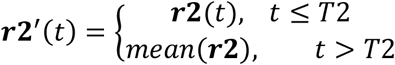

We then obtained the maximal length, Tmax between T2 and T1*tsf:

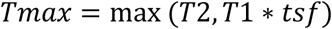

We next compute the mean Euclidian distance d(ssf,tsf) between **r1**^warp^(t) and **r2’**(t) for each pair of ssf and tsf as:

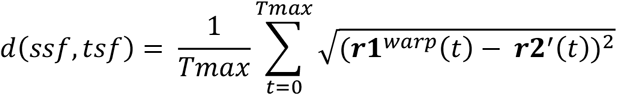

Then the SSF and TSF were defined as the tsf and ssf that gives the minimal d(ssf,tsf):

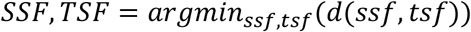

Finally, we defined the SSI as:

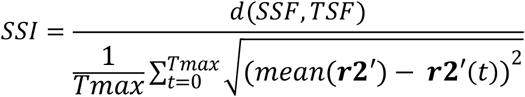

Intuitively, SSI provides a measure of how well the relation from **r1** to **r2** can be explained by temporal and spatial scaling profile, namely the smaller the SSI, the better **r2** can be fitted by warping **r1** temporally and spatially.

### Velocity drive analysis

To understand the transitions between trajectories at different α levels we started from the trajectories at α = 0.9 (**s1)** and α = 0.8 (**s2**)(**Fig 4a**). For the spatial scaling task, **s1** and **s2** naturally have the same length, while for the temporal scaling task, we uniformly subsampled the longer one to the same length as the short one to ensure **s1** and **s2** have the same length. For a given time point on **s2** and its corresponding time point on **s1** with direction **p12** from **s2** to **s1**, there were velocity vectors **v2** and **v1** respectively. Generally, **v2** can be decomposed into the recurrent component, **rec2**, and decay component **d2**. we then sought to compute the angle between **rec2** and **p12** or the angle between **rec2** and **v1** at each corresponding time point on **s1** and **s2**. Finally mean angle across time was obtained for comparison.

### Subspace angle analysis

In **Fig. 4d-f**, we computed the angle between the subspace of the recurrent dynamics at different α levels with the subspace of the output. Specifically, for a given trajectory **r**, we performed the PCA analysis, then the recurrent space was expanded by the first n principal components. The output space was expanded by the learned output weights which led to a 2-dimensional space. Finally, the angle between the recurrent space and output space is computed by the Matlab function *subspace()* between these two spaces.

### RNNs without STP

To study whether STP affects the training and generalization in the temporal or spatial scaling tasks (**Fig. 5**), we modified the standard congruent temporal or spatial scaling task by removing the STP dynamics during training and testing. Specifically, we trained and tested RNNs with x = 1 (equation 2) and u = αU (equation 3) during the whole trial, and other conditions were the same as the standard temporal or spatial scaling task.

### RNNs with an Input-cued-scale approach

To compare the neuromodulation of STP strategy to the strategy of using the input magnitude of an input to cue different scales (**Extended Data Fig. 8**) 23: 1) we removed STP by fixing the variable u at 0.85*U across whole trials; and 2) added an extra input continuously presented across whole trials, the magnitude of which, cued either the length of the trials in the temporal scaling task or the size of the digit in the spatial scaling task. Specifically, 0.9/0.8 corresponded to either 1/1.5 s or 1.5x/1x size respectively. Generalization performance was tested similarly to the α-cued-scale model. To study the natural generalization of the input-cued-scale model, we also trained the RNNs with a single input magnitude level and tested them with different novel levels.

### Simulations of the experimental sensorimotor timing task

To test the potential that modulating α as a universal mechanism for controlling temporal and spatial scale (**Fig. 7**), we simulated two experimental studies of flexible sensorimotor timing tasks (Remington et al., 2018; Soares et al., 2016).

### IAFC task

Same as the experimental conditions on rats for the interval-alternative-forced-choice (IAFC) task (Soares et al., 2016), RNNs were composed of one input for delivering two stimuli lasting 150 ms with a range of intervals, 0.6, 1.05, 1.26,1.38,1.62,1.74,1.95, 2.4 s (short if interval < 1.5 s and long if > 1.5 s) and two outputs corresponding to short or long intervals respectively. The target of the output corresponding to the input interval was set to 1 for a response period of 200 ms right after the second stimulus offset and zeros elsewhere; the other output target was zeros everywhere. The decision was made based on the mean activity during the response period for the two outputs in a winner-take-all manner and performance was defined as the percentage of the correct decision trials. RNNs were trained with α = 0.8 for all units and tested with 0.9 and to simulate the effect of optogenetic inactivation or activation of dopamine activity respectively. We set σ (in equation 1) to 1 to match the noise level of the experiments. Same as the experiment, we fitted the long choice probability by a sigmoid function:

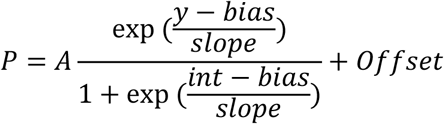

where y is the input intervals.

### Flexible sensorimotor timing task

For the simulation of the Ready-Set-Go task (Remington et al., 2018), the input of the RNN delivered two stimuli lasting 100 ms with the interval from the pool of 7 intervals uniformly spaced in 0.5-1s (sensory time, t_s_). Based on the context cued by α = 0.9 or α = 0.8, the output unit should generate a linear ramp (0-1) crossing threshold (0.75) at 1× or 1.5× as input intervals (the target time t_t_) since the offset of the second stimulus respectively (production time, tp). Same as the experiments, we defined one trial with tp as correct if the error0020= |t_p_ -t_t_| was smaller than 0.2* t_p_ + 0.025 s. Again performance was defined as the percentage of the correct trials.

## Data and codes availability

All data are available in the main text or supplementary materials. Codes used for the simulations in this paper will be available at (https://github.com/ShanglinZhou/Temporal_Spatial_Scale_STP).

## Supporting information

**Extended Data Fig. 1:**
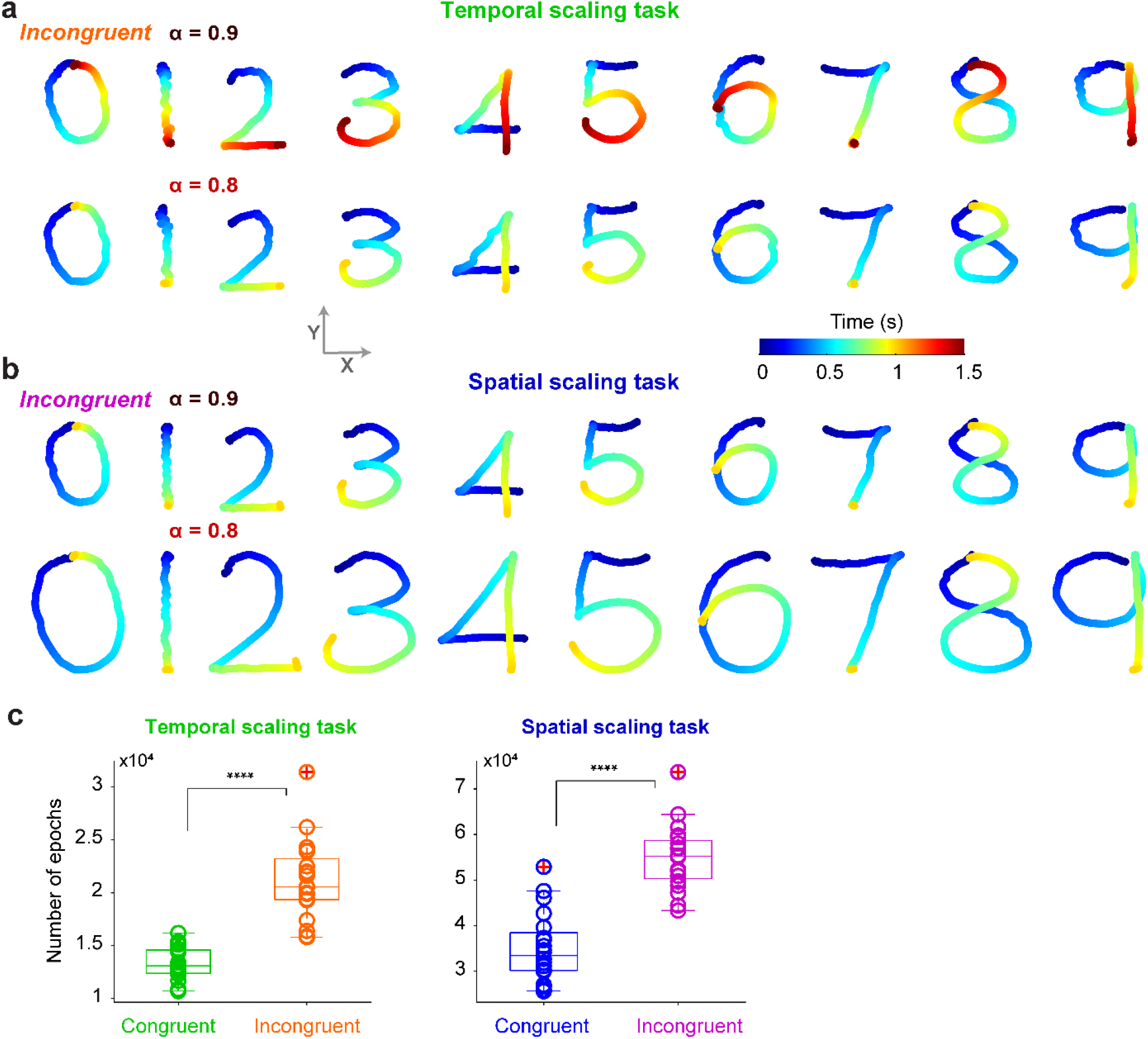
Example output traces for the incongruent cases in temporal and spatial tasks. **a**, Output traces of an example RNN at α = 0.9 (top) and α = 0.8 (bottom) for the incongruent cases in temporal scaling task. **b**, Same as **a** but in spatial scaling task.). **c**, Comparison of the number of training epochs for RNNs for congruent and incongruent settings in temporal scaling task (left) and spatial scaling task right (n = 20 RNNs; P < 10^−7^, P < 10^−6^, two-sided Wilcoxon rank sum test respectivly). Boxplot: central lines, median; bottom and top edges, lower and upper quartiles; whiskers, extremes; red cross, outliers.

**Extended Data Fig. 2:**
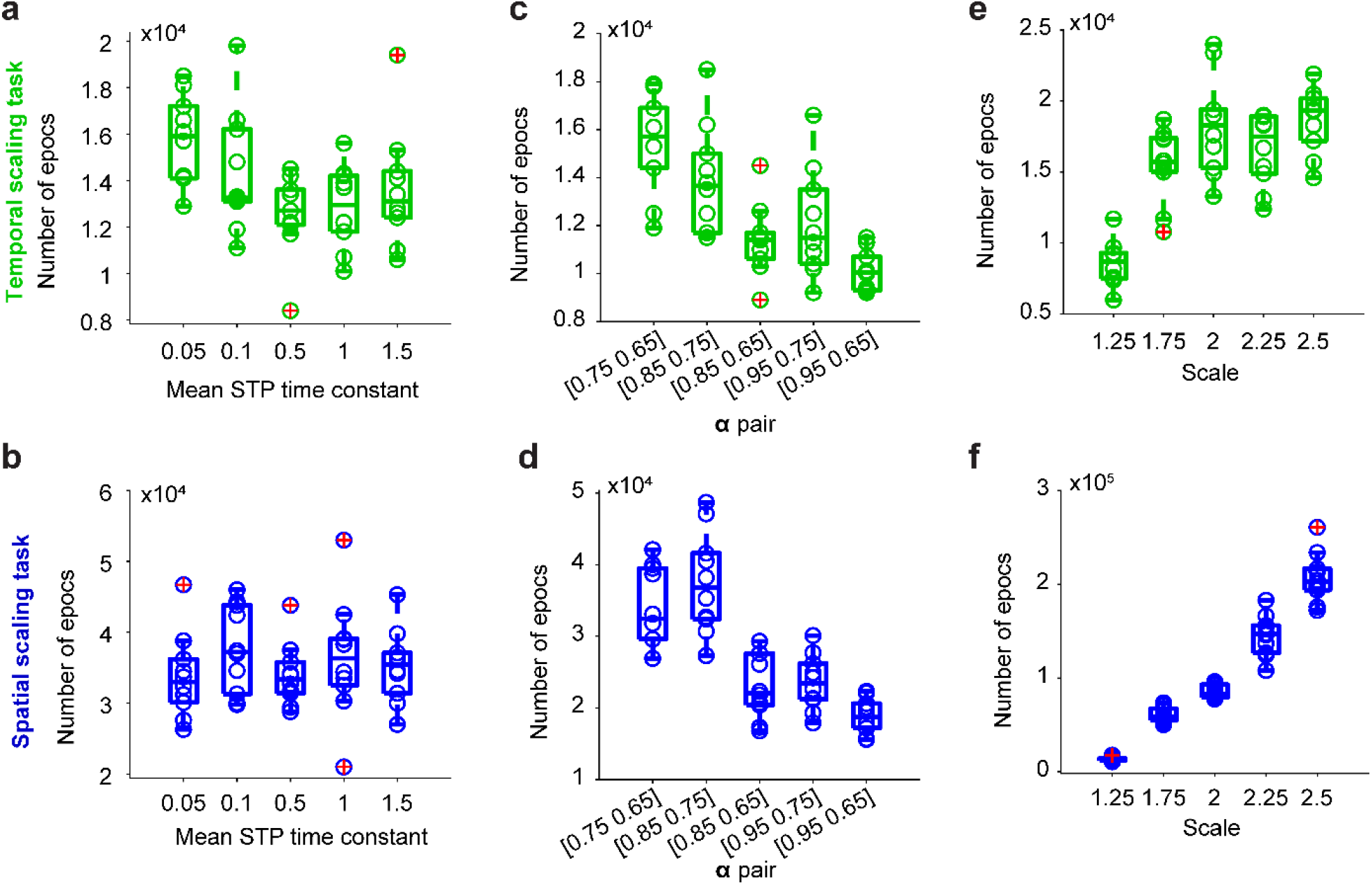
Learning temporal and spatial scaling tasks is robust across a diverse range of hyperparameters. The number of training epochs needed to reach the same success criterion in temporal (top) and spatial (bottom) scaling tasks for a diverse range of hyperparameters: mean of τd and τx in the STP model (**a**,**b**), pair of α level used for different scales (**c**,**d**), and scale factor between the two scales (**e**,**f**).

**Extended Data Fig. 3:**
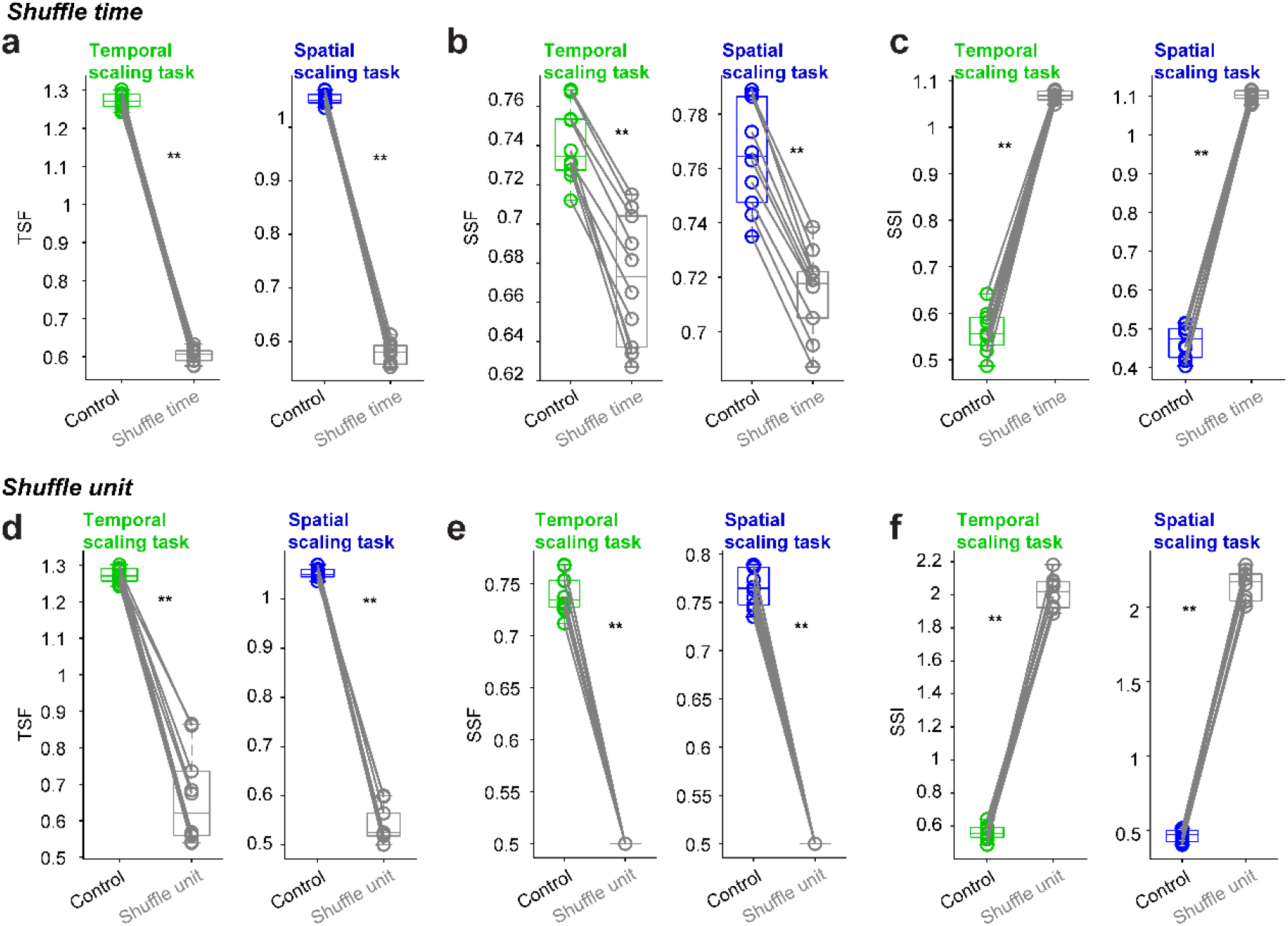
Shuffling time or units both disrupted the temporal-spatial profile of recurrent dynamics. **a**, comparison of average TSF across 20 RNNs between control and time-shuffled congruent case in temporal (left) and spatial (right) scaling task. **b**, Same as a but for SSF. c, Same as a but for SSI. d,e,f, Same as a,b,c but for shuffling units but keeping the temporal structure of each unit. In all the conditions, shuffling either time or units significantly disrupted the temporal-spatial profile (n = 10 digits; P = 0.002, two-sided Wilcoxon signed rank test for all conditions).

**Extended Data Fig. 4:**
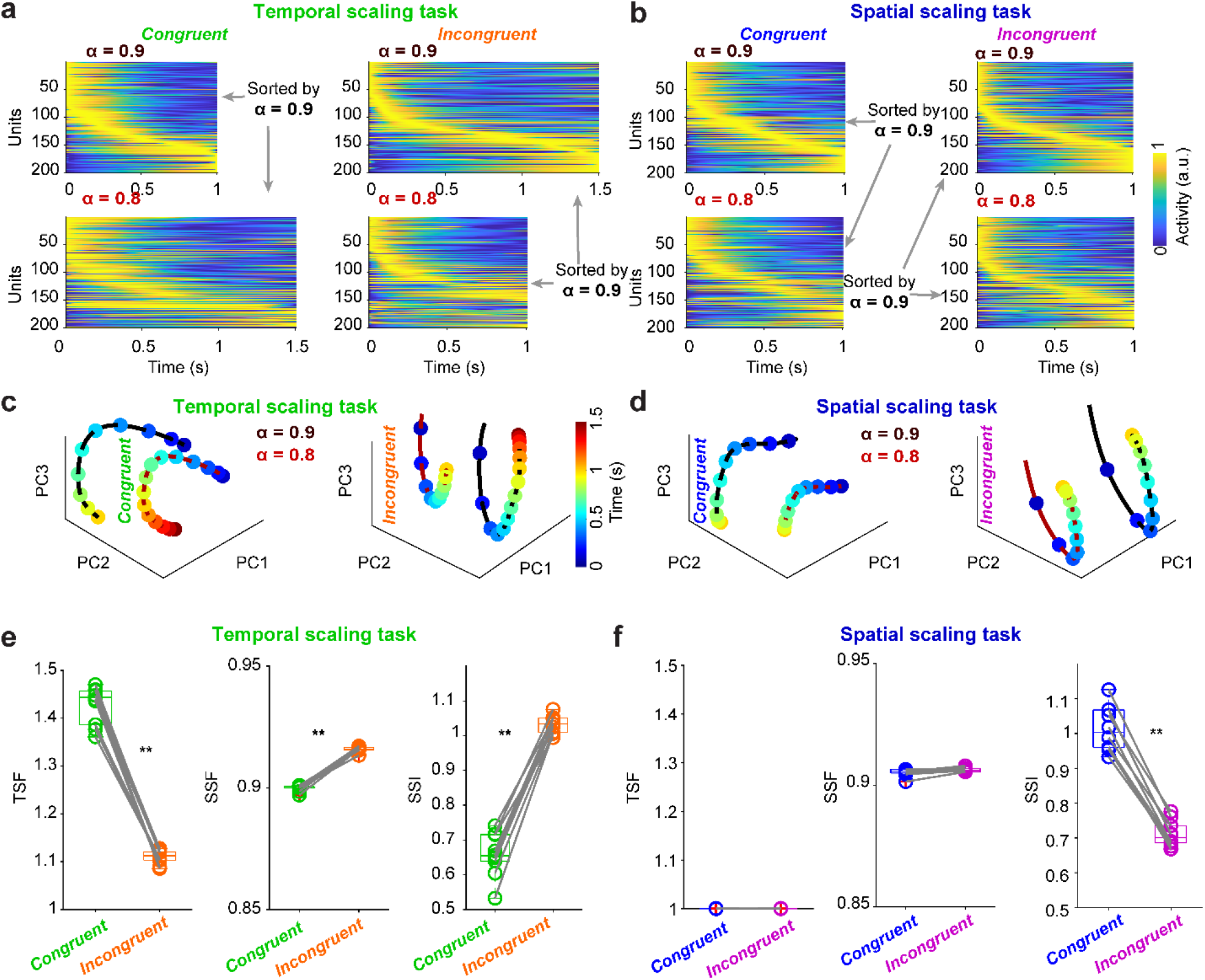
Synaptic efficacy dynamics exhibited a similar temporal-spatial profile as the activity dynamics. **a**, Normalized synaptic efficacy (x*u in STP) at α = 0.9 (top) and α = 0.8 (bottom) sorted according to the peak latency at α = 0.9 for congruent (left) and incongruent (right) temporal scaling task. The Red dashed line denoted the time point of 0.5 s. **b**, Same as **a** but for spatial scaling task. **c**, Plot of the first three principal components of synaptic efficacy of α = 0.9 (black) and α = 0.8 (dark red) for congruent (left) and incongruent (right) cases in temporal scaling task. Color codes the time. **d**, Same as c but for spatial scaling task. **e**, comparison of congruent and incongruent cases for average TSF across 20 RNNs in the temporal (left) and spatial (right) scaling task (n = 10 digits; P = 0.002 and 1, two-sided Wilcoxon signed rank test for temporal and spatial scaling task respectively). **f**, Same as **e** but for SSF (n = 10 digits; P = 0.002 and 0.152, two-sided Wilcoxon signed rank test for temporal and spatial scaling task respectively). **g**, Same as **e** but for SSI (n = 10 digits; P = 0.002, two-sided Wilcoxon signed rank test for both temporal and spatial scaling task). Boxplot: central lines, median; bottom and top edges, lower and upper quartiles; whiskers, extremes; red cross, outliers.

**Extended Data Fig. 5:**
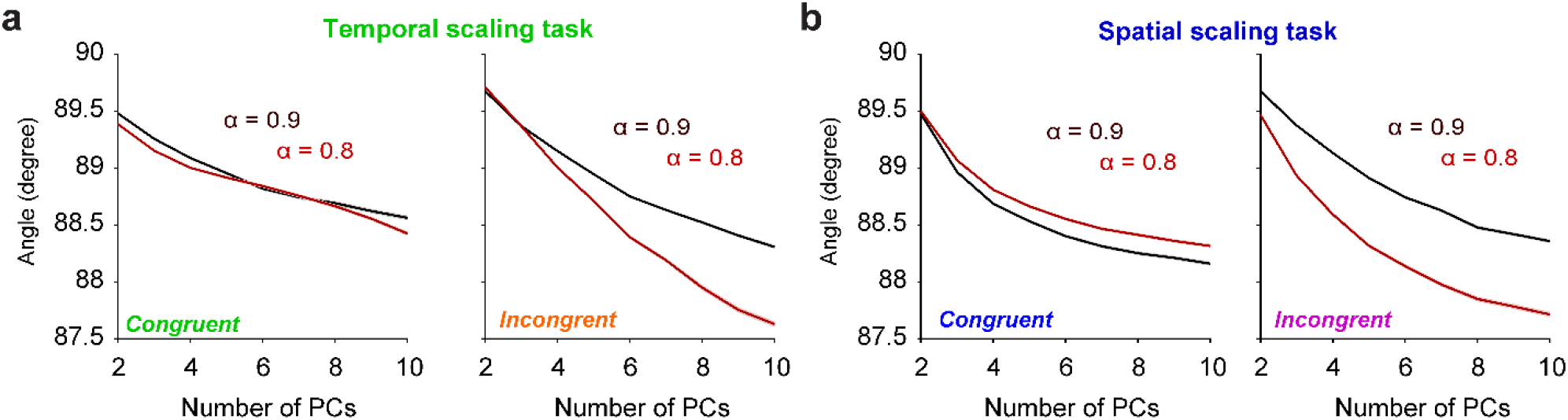
Angle between recurrent and output space is robust across a diverse range of recurrent PC numbers. **a**, Average angle between output space and recurrent space expanded by different number of PCs at α = 0.9 (black) and 0.8 (red) for congruent (left) and incongruent (right) cases in temporal scaling task. **b**, Same as **a** but for spatial scaling task.

**Extended Data Fig. 6:**
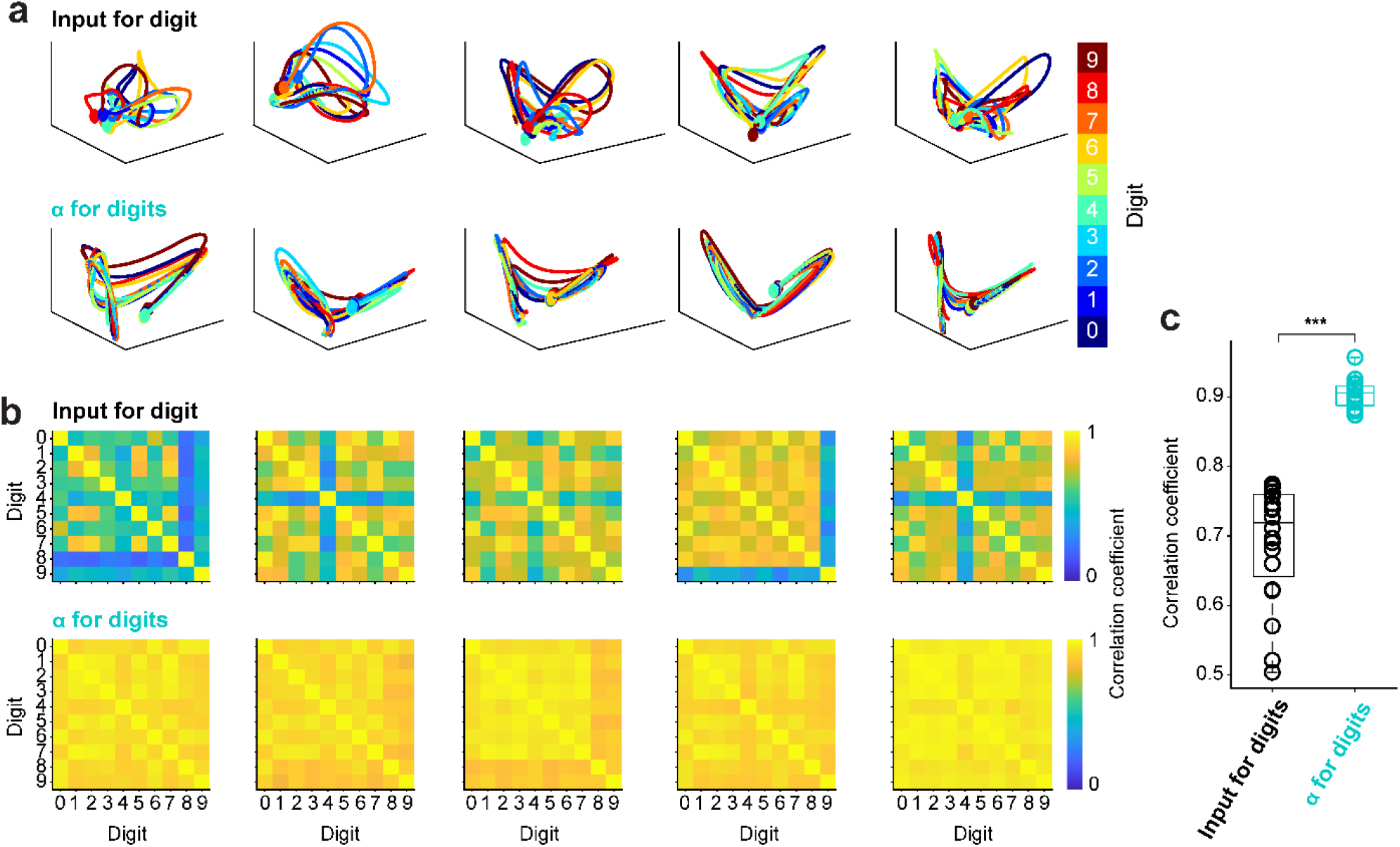
Different dynamical profiles between RNNs with input or α signaling digits in joint temporal-spatial scaling task. **a**, PCA plot of the recurrent dynamics in five example RNNs with input (top) and α (bottom) signaling digit. Color codes digits. **b**, Cross-digit population correlation in the five example RNNs in **a** with input (top) and α (bottom) signaling digit. **c**, Comparison of the average cross-digit correlation in **b** between RNNs with input and α signaling digits (n = 20 RNNs; P < 10^−7^, two-sided Wilcoxon rank sum test). Boxplot: central lines, median; bottom and top edges, lower and upper quartiles; whiskers, extremes; red cross, outliers.

**Extended Data Fig. 7:**
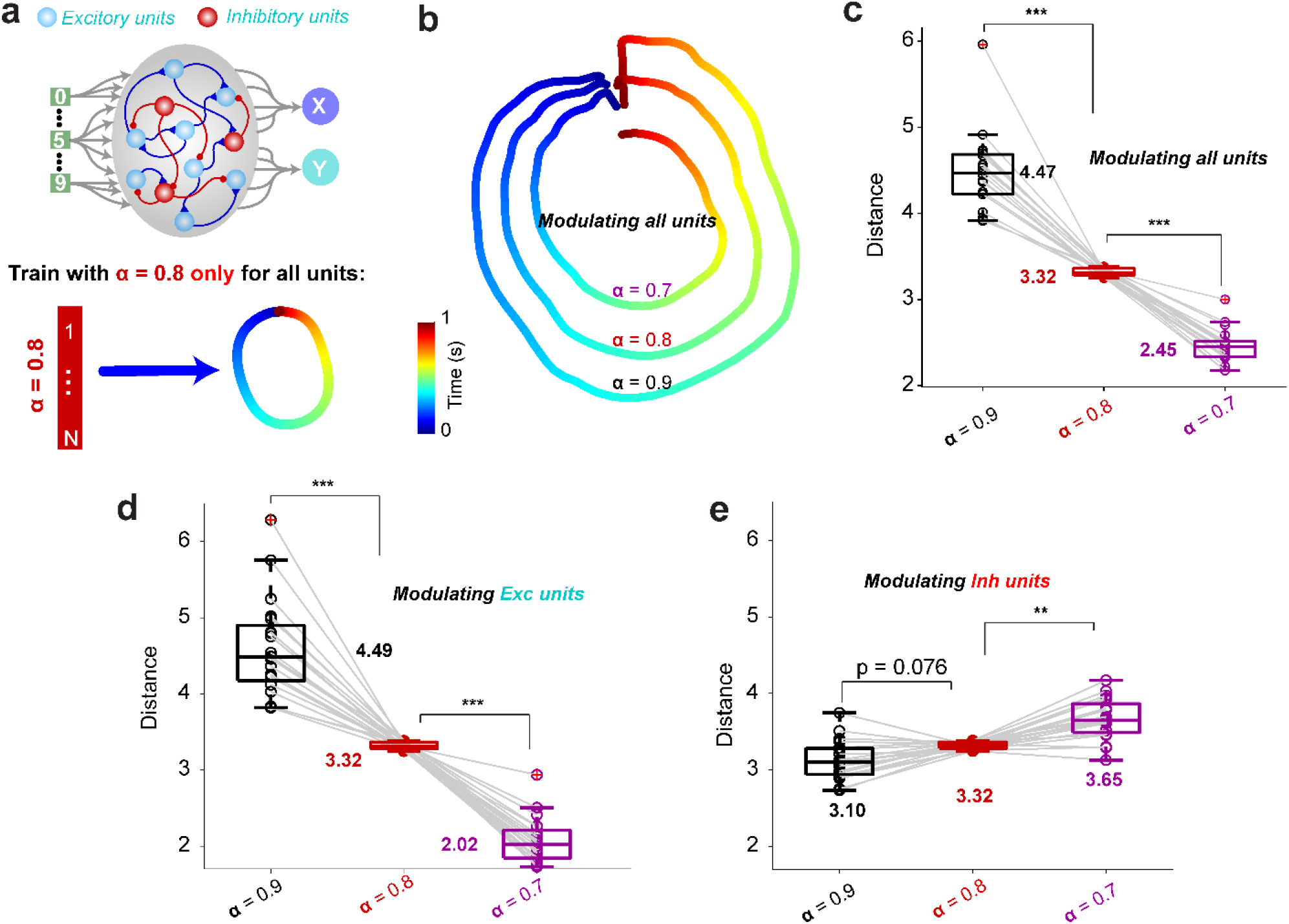
RNNs trained with a single α level generalize congruently to novel levels. **a**, Schematic of RNNs trained with single α = 0.8 for all units with a target of the short-small digit. **b**, Example output traces of digit 0 at α = 0.7 and 0.9 for all units with α = 0.8 shown for comparison. **c**, Summary of the distance for different α. Decreasing α significantly decreases the distance (n = 20 RNNs, Kruskal-Wallis test, P<10^−11^, χ^2^_(2,57)_ = 52.5) and distance for α = 0.9 and 0.7 is significantly higher and lower than that for α = 0.8 respectively (P = 0.0009 for both, by Dunn’s multiple comparison test). **d**, Same as **c** but for modulating α only for excitatory units. Decreasing α significantly decrease the distance (n = 20 RNNs, Kruskal-Wallis test, P<10^−11^, χ^2^_(2,57)_ = 52.5) and distance for α = 0.9 and 0.7 is significantly higher and lower than that for α = 0.8 respectively (P = 0.0009 for both, by Dunn’s multiple comparison test). **e**, Same as **c** but for modulating α only for inhibitory units. Decreasing α significantly increases the distance (n = 20 RNNs, Kruskal-Wallis test, P<10^61^, χ^2^_(2,57)_ = 31.1) and distance for α = 0.9 and 0.7 is lower and higher than that for α = 0.8 respectively (P = 0.076 and 0.003 respectively by Dunn’s multiple comparison test).

**Extended Data Fig. 8:**
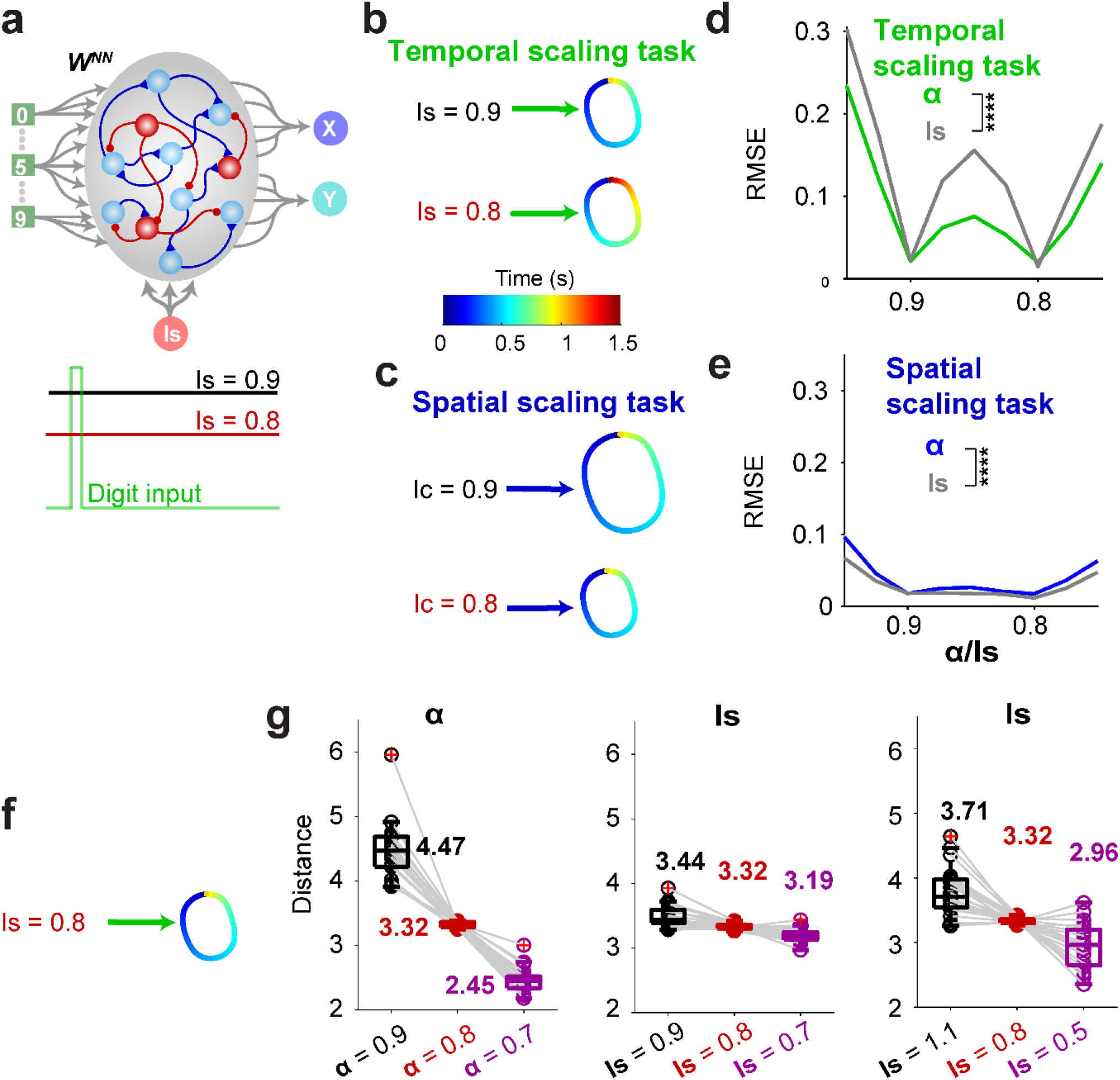
RNNs trained with input magnitude cueing the scales. **a**, Schematic of RNNs trained with an extra input Is continusely presented during the whole trial to cue either the temporal or spatial scales. **b**,Simialr to the congruent settings for the standard α-cued-scale approach, the ls = 0.9/0.8 correspond to the short/long duration respectively. **c**, Simialr to **b** but for the spatial scaling task. **d**, Comparision of the generalization performance for the two approach: input-cued-scale (gray) and α-cued-scale (green) (n = 20 RNNs; two-way ANOVA with mixed-effect design, F_1,38_ = 885.2, P < 10^-27^). **e**, Same as d but for the spatial scaling tasks (n = 20 RNNs; two-way ANOVA with mixed-effect design, F_1,38_ = 47.1, P < 10^-7^). **f**, Schematic of RNN with input-cued-scale approach but trained with single input magnitude level (Is = 0.8). **g**, The output distance of RNNs trained as **f** and tested with Is = 0.9 and 0.7 (midlle) or with Is = 1.1 and 0.5 for a wider range (right). Resutls for the α-cued-scale approach are presented for comparison (left). The distance median are shown on top of each codintions.

## REFERENCES

Abbott, L.F., and Regehr, W.G. (2004). Synaptic computation. Nature 431, 796–803.

Baimoukhametova, D.V., Hewitt, S.A., Sank, C.A., and Bains, J.S. (2004). Dopamine Modulates Use-Dependent Plasticity of Inhibitory Synapses. The Journal of Neuroscience 24, 5162–5171.

Barak, O., and Tsodyks, M. (2014). Working models of working memory. Current Opinion in Neurobiology 25, 20–24.

Buhusi, C.V., and Meck, W.H. (2002). Differential effects of methamphetamine and haloperidol on the control of an internal clock. Behav Neurosci 116, 291–297.

Buonomano, D.V., and Merzenich, M.M. (1995). Temporal information transformed into a spatial code by a neural network with realistic properties. Science 267, 1028–1030.

Burke, K.J., Jr., Keeshen, C.M., and Bender, K.J. (2018). Two Forms of Synaptic Depression Produced by Differential Neuromodulation of Presynaptic Calcium Channels. Neuron 99, 969-984.e967.

Chaisangmongkon, W., Swaminathan, S.K., Freedman, D.J., and Wang, X.-J. (2017). Computing by Robust Transience: How the Fronto-Parietal Network Performs Sequential, Category-Based Decisions. Neuron 93, 1504-1517.e1504.

Chiu, C.Q., Puente, N., Grandes, P., and Castillo, P.E. (2010). Dopaminergic Modulation of Endocannabinoid-Mediated Plasticity at GABAergic Synapses in the Prefrontal Cortex. Journal of Neuroscience 30, 7236–7248.

Churchland, M.M., Cunningham, J.P., Kaufman, M.T., Foster, J.D., Nuyujukian, P., Ryu, S.I., and Shenoy, K.V. (2012). Neural population dynamics during reaching. Nature 487, 51–56.

Cicchini, G.M., Arrighi, R., Cecchetti, L., Giusti, M., and Burr, D.C. (2012). Optimal Encoding of Interval Timing in Expert Percussionists. The Journal of Neuroscience 32, 1056–1060.

Crowe, D.A., Zarco, W., Bartolo, R., and Merchant, H. (2014). Dynamic Representation of the Temporal and Sequential Structure of Rhythmic Movements in the Primate Medial Premotor Cortex. The Journal of Neuroscience 34, 11972–11983.

Drew, M.R., Fairhurstb, S., Malapani, C., Horvitz, J.C., and Balsam, P.D. (2003). Effects of dopamine antagonists on the timing of two intervals. Pharmacology, Biochemistry and Behavior 75, 9–15.

Fung, B.J., Sutlief, E., and Hussain Shuler, M.G. (2021). Dopamine and the interdependency of time perception and reward. Neuroscience & Biobehavioral Reviews 125, 380–391.

Gao, W.J., Krimer, L.S., and Goldman-Rakic, P.S. (2001). Presynaptic regulation of recurrent excitation by D1 receptors in prefrontal circuits. P Natl Acad Sci USA 98, 295–300.

Gao, W.J., Wang, Y., and Goldman-Rakic, P.S. (2003). Dopamine modulation of perisomatic and peridendritic inhibition in prefrontal cortex. Journal of Neuroscience 23, 1622–1630.

Gonzalez-Islas, C., and Hablitz, J.J. (2003). Dopamine Enhances EPSCs in Layer II–III Pyramidal Neurons in Rat Prefrontal Cortex. The Journal of Neuroscience 23, 867–875.

Goudar, V., and Buonomano, D.V. (2018). Encoding sensory and motor patterns as time-invariant trajectories in recurrent neural networks. Elife 7.

Hardy, N.F., Goudar, V., Romero-Sosa, J.L., and Buonomano, D.V. (2018). A model of temporal scaling correctly predicts that motor timing improves with speed. Nature Communications 9, 4732.

Harpaz, N.K., Flash, T., and Dinstein, I. (2014). Scale-Invariant Movement Encoding in the Human Motor System. Neuron 81, 452–462.

Hennequin, G., Vogels Tim P., and Gerstner, W. (2014). Optimal Control of Transient Dynamics in Balanced Networks Supports Generation of Complex Movements. Neuron 82, 1394–1406.

Kim, R., Li, Y., and Sejnowski, T.J. (2019). Simple framework for constructing functional spiking recurrent neural networks. Proc Natl Acad Sci U S A 116, 22811–22820.

Kroener, S., Chandler, L.J., Phillips, P.E.M., and Seamans, J.K. (2009). Dopamine Modulates Persistent Synaptic Activity and Enhances the Signal-to-Noise Ratio in the Prefrontal Cortex. PLoS ONE 4, e6507.

Laje, R., and Buonomano, D.V. (2013). Robust timing and motor patterns by taming chaos in recurrent neural networks. Nat Neurosci 16, 925–933.

Lake, J.I., and Meck, W.H. (2013). Differential effects of amphetamine and haloperidol on temporal reproduction: Dopaminergic regulation of attention and clock speed. Neuropsychologia 51, 284–292.

Leyrer-Jackson, J.M., and Thomas, M.P. (2018). Layer-specific effects of dopaminergic D1 receptor activation on excitatory synaptic trains in layer V mouse prefrontal cortical pyramidal cells. Physiological Reports 6, e13806.

Lindén, H., Petersen, P.C., Vestergaard, M., and Berg, R.W. (2022). Movement is governed by rotational neural dynamics in spinal motor networks. Nature.

Mante, V., Sussillo, D., Shenoy, K.V., and Newsome, W.T. (2013). Context-dependent computation by recurrent dynamics in prefrontal cortex. Nature 503, 78–84.

Maricq, A.V., and Church, R.M. (1983). The differential effects of haloperidol and methamphetamine on time estimation in the rat. Psychopharmacology (Berl) 79, 10–15.

Markram, H., Wang, Y., and Tsodyks, M. (1998). Differential signaling via the same axon of neocortical pyramidal neurons. Proc Natl Acad Sci USA 95, 5323–5328.

Masse, N.Y., Yang, G.R., Song, H.F., Wang, X.J., and Freedman, D.J. (2019). Circuit mechanisms for the maintenance and manipulation of information in working memory. Nature Neuroscience 22, 1159-+.

Meck, W.H. (1996). Neuropharmacology of timing and time perception. Cog Brain Res 3, 227–242.

Merchant, H., Pérez, O., Bartolo, R., Méndez, J.C., Mendoza, G., Gámez, J., Yc, K., and Prado, L. (2015). Sensorimotor neural dynamics during isochronous tapping in the medial premotor cortex of the macaque. European Journal of Neuroscience 41, 586–602.

Mongillo, G., Barak, O., and Tsodyks, M. (2008). Synaptic Theory of Working Memory. Science 319, 1543–1546.

Motanis, H., Seay, M.J., and Buonomano, D.V. (2018). Short-Term Synaptic Plasticity as a Mechanism for Sensory Timing. Trends in neurosciences 41, 701–711.

Murray, J.D., Bernacchia, A., Roy, N.A., Constantinidis, C., Romo, R., and Wang, X.-J. (2017). Stable population coding for working memory coexists with heterogeneous neural dynamics in prefrontal cortex. Proceedings of the National Academy of Sciences 114, 394–399.

Murray, J.M., and Escola, G.S. (2017). Learning multiple variable-speed sequences in striatum via cortical tutoring. eLife 6, e26084.

Nadim, F., and Bucher, D. (2014). Neuromodulation of neurons and synapses. Current Opinion in Neurobiology 29, 48–56.

Panigrahi, B., Martin, K.A., Li, Y., Graves, A.R., Vollmer, A., Olson, L., Mensh, B.D., Karpova, A.Y., and Dudman, J.T. (2015). Dopamine Is Required for the Neural Representation and Control of Movement Vigor. Cell 162, 1418–1430.

Rammsayer, T.H. (1999). Neuropharmacological evidence for different timing mechanisms in humans. Q J Exp Psychol B 52, 273–286.

Remington, E.D., Narain, D., Hosseini, E.A., and Jazayeri, M. (2018). Flexible Sensorimotor Computations through Rapid Reconfiguration of Cortical Dynamics. Neuron 98, 1005-1019.e1005.

Rosenbaum, D.A. (2010). Human motor control, 2nd edn (Amsterdam; Boston, MA: Elsevier Inc).

Rush, A.M., Kilbride, J., Rowan, M.J., and Anwyl, R. (2002). Presynaptic group III mGluR modulation of short-term plasticity in the lateral perforant path of the dentate gyrus in vitro. Brain Research 952, 38–43.

Saxena, S., Russo, A.A., Cunningham, J., and Churchland, M.M. (2022). Motor cortex activity across movement speeds is predicted by network-level strategies for generating muscle activity. Elife 11.

Schultz, W., Dayan, P., and Montague, P.R. (1997). A neural substrate of prediction and reward. Science 275, 1593–1599.

Seamans, J.K., Durstewitz, D., Christie, B.R., Stevens, C.F., and Sejnowski, T.J. (2001a). Dopamine D1/D5 receptor modulation of excitatory synaptic inputs to layer V prefrontal cortex neurons. Proc Natl Acad Sci U S A 98, 301–306.

Seamans, J.K., Gorelova, N., Durstewitz, D., and Yang, C.R. (2001b). Bidirectional dopamine modulation of GABAergic inhibition in prefrontal cortical pyramidal neurons. Journal of Neuroscience 21, 3628–3638.

Simen, P., and Matell, M. (2016). Why does time seem to fly when we’re having fun? Science 354, 1231–1232.

Soares, S., Atallah, B.V., and Paton, J.J. (2016). Midbrain dopamine neurons control judgment of time. Science 354, 1273–1277.

Stroud, J.P., Porter, M.A., Hennequin, G., and Vogels, T.P. (2018). Motor primitives in space and time via targeted gain modulation in cortical networks. Nature Neuroscience 21, 1774–1783.

Sussillo, D., and Abbott, L.F. (2009). Generating Coherent Patterns of Activity from Chaotic Neural Networks. Neuron 63, 544–557.

Tecuapetla, F., Carrillo-Reid, L., Bargas, J., and Galarraga, E. (2007). Dopaminergic modulation of short-term synaptic plasticity at striatal inhibitory synapses. Proceedings of the National Academy of Sciences 104, 10258–10263.

Tritsch, N.X., and Sabatini, B.L. (2012). Dopaminergic Modulation of Synaptic Transmission in Cortex and Striatum. Neuron 76, 33–50.

Tsodyks, M.V., and Markram, H. (1997). The neural code between neocortical pyramidal neurons depends on neurotransmitter release probability. P Natl Acad Sci USA 94, 719–723.

Vyas, S., Golub, M.D., Sussillo, D., and Shenoy, K.V. (2020). Computation Through Neural Population Dynamics. Annual Review of Neuroscience 43, 249–275.

Wang, J., Narain, D., Hosseini, E.A., and Jazayeri, M. (2018). Flexible timing by temporal scaling of cortical responses. Nature Neuroscience 21, 102–110.

Zhou, S., Masmanidis, S.C., and Buonomano, D.V. (2020). Neural Sequences as an Optimal Dynamical Regime for the Readout of Time. Neuron 108, 651-+.

Zhou, S., Masmanidis, S.C., and Buonomano, D.V. (2022). Encoding time in neural dynamic regimes with distinct computational tradeoffs. Plos Computational Biology 18.

Zucker, R.S., and Regehr, W.G. (2002). Short-term synaptic plasticity. Annual review of physiology 64, 355–405.

